# Comparative transcriptomics reveals shared and context-dependent adaptations of bovine cells across culture systems for cultivated meat

**DOI:** 10.64898/2026.11.18.767523

**Authors:** Ivana Ribarski-Chorev, Moritz Burghardt, Shadi Tawil, Alon Gershkoviz, Dafna Shpatz, Gisele Schudy, Yarden Beh Hamo, Yael Gilad, Ilana Vainshtein, Adar Kfir, Tamir Tuller, Sharon Schlesinger

**Affiliations:** Department of Animal Sciences, The Robert H. Smith Faculty of Agriculture, Food and Environment, The Hebrew University of Jerusalem, Rehovot, 7610001, Israel; Department of Biomedical Engineering, Tel Aviv University, Tel Aviv-Yafo, 6997801, Israel; Sagol School of Neuroscience, Tel Aviv University, Tel Aviv-Yafo, 6997801, Israel

## Abstract

Developing cell-culture systems for cultivated meat requires understanding how candidate cells respond to prolonged expansion and different growth environments. However, the extent to which bovine cells share transcriptional adaptations across these conditions, or retain responses specific to their origin and cellular state, remains unclear. Here, we systematically profiled bovine umbilical cord (BUC)-and adipose tissue (BAT)-derived mesenchymal stem cells (MSCs) across different culture architectures, passage stages, and differentiation states, including immortalized BUC cells and satellite cells as a myogenic reference.

We show that global transcriptomic variation is strongly associated with culture architecture, cell identity, and cellular state. Immortalization preserved several early-passage transcriptional features, particularly proliferation-related programs, while opposing selected changes associated with primary cell aging. Culture architecture induced distinct rather than uniform three-dimensional responses. Spheroid formation produced a pronounced transcriptional shift characterized by reduced proliferation-, chromatin-, and development-associated programs, alongside changes in extracellular-matrix organization, metabolism, and intracellular remodeling. Across primary umbilical cord-derived, adipose-derived, and myogenically induced umbilical cord-derived cells, scaffold-versus-plate comparisons revealed correlated but non-identical transcriptional responses. These involved changes in extracellular-matrix organization, adhesion, membrane trafficking, and developmental pathways, with endocytosis enriched across all three comparisons among the pathways examined. During myogenic induction, both satellite cells and BUC-derived cells exhibited muscle-related cytoskeletal and contractile programs, while scaffold culture further supported myogenic signatures in BUC-derived cells.

These findings distinguish recurring culture-associated responses from transcriptional features tied to cell source and expansion history. They support evaluating cultivated-meat candidate cells in their intended culture systems, not solely under conventional plate conditions.

## Introduction

The development of cultivated meat still faces major bottlenecks, particularly the need to expand cells efficiently at high density, direct them reproducibly toward desired lineages, and reduce the cost and complexity of culture systems and media (Post et al., 2020). Among the candidate cell types for these applications, mesenchymal stromal/stem cells (MSCs) are especially promising because they can be isolated from accessible bovine tissues, expanded in culture, and directed toward mesenchymal fates relevant to muscle and fat production. In culture, MSCs are classically defined by their fibroblast-like morphology, characteristic marker profile, and capacity for tri-lineage differentiation into osteogenic, adipogenic, and chondrogenic lineages (Dominici et al., 2006; Keating, 2012). At the same time, cell fate, proliferation, and functional properties are strongly influenced by the physical and environmental context, including substrate properties, spatial organization, and other culture-associated stresses (Engler et al., 2006; Ferreira et al., 2013; Hu and Li, 2018; Phelps et al., 2023; Pittenger et al., 2019). This question is especially important for cultivated meat production, where the key challenge is not only whether cells grow, but whether they do so reproducibly across culture architectures that are compatible with scale-up and robust manufacturing (Hauser et al., 2024; Klatt et al., 2024).

Previous work from our lab established a panel of bovine cell systems from distinct tissue sources and culture configurations, including primary MSCs derived from bovine umbilical cord (BUC) and bovine adipose tissue (BAT), immortalized umbilical cord-derived MSCs, myogenically induced MSCs, and bovine satellite cells in both primary and differentiated states (Tawil et al., 2026; Gershkoviz et al., 2026; Ribarski-Chorev et al., 2023; Gome et al., 2024; Shimoni et al., 2020). These cells were grown under multiple spatial conditions, including conventional tissue-culture plates, porous cellulose-based scaffolds, and spheroid culture, and were profiled by RNA-seq across different cellular states and passages (Figure 1, Supplementary Table S1). Together, these datasets provide a unique opportunity to compare, within a unified analytical framework, the effects of tissue origin, immortalization, passage-associated changes, three-dimensional organization, and myogenic differentiation on bovine stem and progenitor cell transcriptional states.

**Figure 1.**
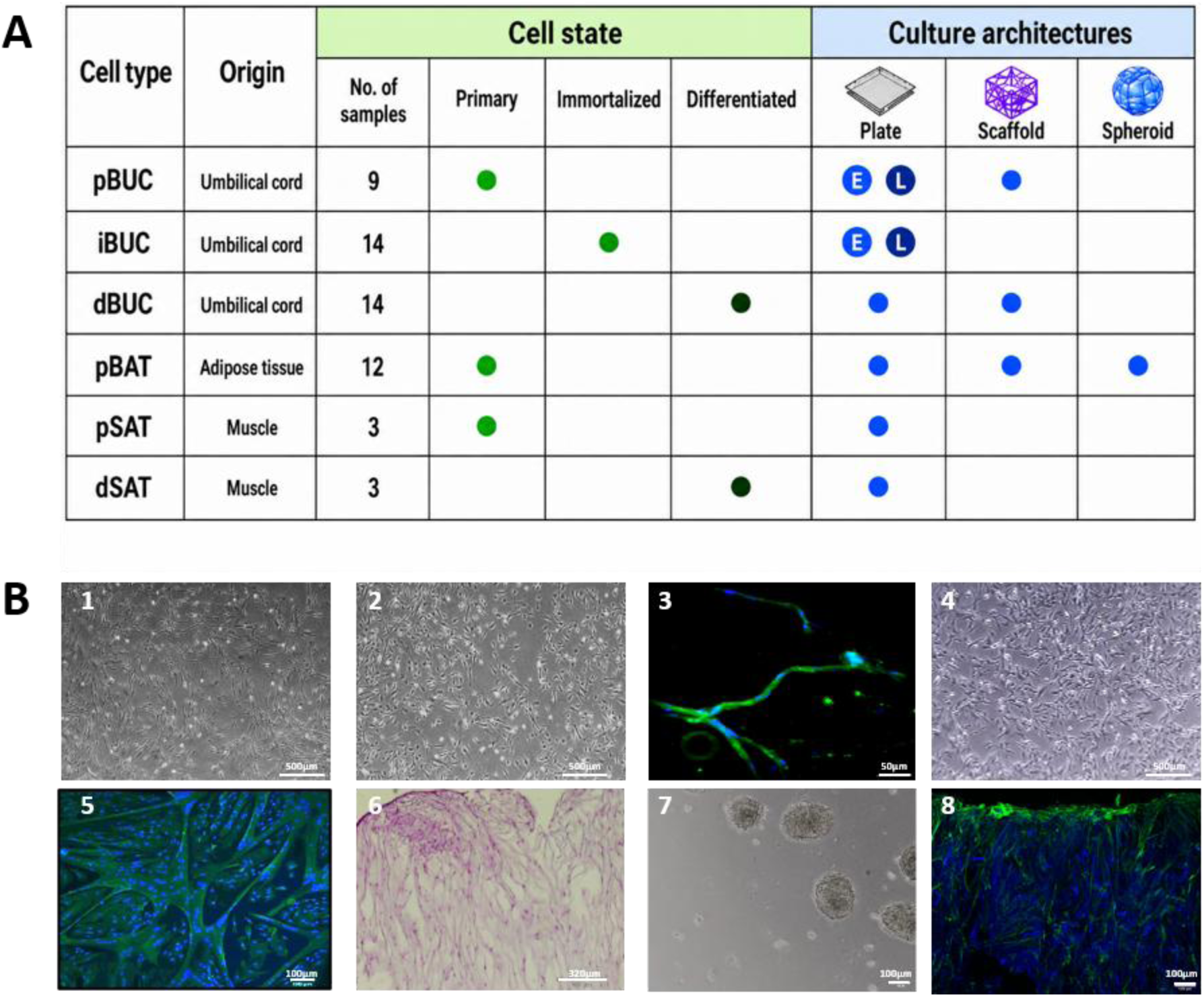
Experimental design of bovine culture systems. **A.** Comparison matrix summarizing the tissue origin, culture architecture, passage group, and cellular state of the samples included in the study. The analyzed cells comprised primary bovine umbilical cord-derived mesenchymal stromal cells (pBUC), immortalized BUC-derived cells (iBUC), myogenically induced BUC-derived cells (dBUC), primary bovine adipose tissue-derived mesenchymal stromal cells (pBAT), primary bovine satellite cells (pSAT), and differentiated bovine satellite cells (dSAT). Culture architecture included conventional two-dimensional plate culture (pl), three-dimensional cellulose scaffold culture (sc), and three-dimensional spheroid (sp) suspension culture. The matrix also indicates the primary or immortalized status, early-and late-passage groups (E = 12 or 21 days and L = 50 or 165 days in culture, respectively), and myogenic induction or differentiation state represented in the dataset. **B.** Representative images of the cell models and culture conditions included in the transcriptomic analysis: pBUC_12pl (B1), pBAT_pl (B2), dBUC_21pl (B3), iBUC_21pl (B4), dSAT_3pl (B5), pBUC_sc (B6), pBAT_35sp (B7), and dBUC_sc (B8). B3, B5, and B8 show immunofluorescence staining for titin (green), with nuclei counterstained using DAPI (blue), and were imaged by confocal microscopy. B6 shows hematoxylin and eosin staining of bovine MSCs cultured for one week on 1% cellulose nanocrystal scaffolds. B1, B2, B4, and B7 are representative bright-field images. Scale bars are shown in the corresponding images.

While several studies have already compared transcriptomes across bovine MSC tissue sources, analyzed transcriptomic remodeling during bovine satellite-cell proliferation and differentiation, or examined scaffold-dependent transcriptional changes in cultured muscle systems, these analyses have generally addressed one variable at a time (Bezjak et al., 2024; Danev et al., 2024; Lu et al., 2024). In the present study, we performed a comprehensive bioinformatic analysis of bulk RNA-seq data generated from these previously isolated and characterized bovine cell populations. Specifically, we compared primary bovine umbilical cord MSCs, primary bovine adipose-derived MSCs, and immortalized bovine umbilical cord MSCs at early and late passages. We also compared myogenically induced MSCs, together with primary and differentiated bovine satellite cells, across plate and scaffold conditions. By jointly analyzing lineage, passage, immortalization, and culture architecture rather than considering each variable separately, we sought to define the shared and distinct transcriptional adaptations associated with cell identity and spatial organization, and to determine how these variables shape proliferation-related programs, extracellular matrix organization, metabolic state, stress responses, and lineage-associated gene expression. To our knowledge, no previous study has systematically compared the transcriptomic effects of tissue origin, immortalization, passage, myogenic induction, and culture architecture across multiple bovine stem and progenitor cell models within a single analytical framework (Hauser et al., 2024).

Using this integrated dataset, we examined how cell identity, tissue origin, passage, immortalization, differentiation state, and culture architecture are associated with global transcriptomic organization and biological-process enrichment. We hypothesized that cell source and cellular state would define the major transcriptional differences among the models, whereas scaffold and spheroid culture would induce architecture-associated responses involving extracellular matrix organization, adhesion, mechanotransduction, metabolism, and lineage-related programs. This comparative framework was designed to distinguish broadly conserved culture responses from transcriptional features specific to individual bovine cell models.

## Results

### Experimental design and transcriptomic dataset overview

To examine how cell identity, cellular state, and culture architecture shape transcriptional profiles, we performed bulk RNA sequencing across a panel of bovine cells comprising primary umbilical cord-derived mesenchymal stromal cells (pBUC), primary adipose tissue-derived mesenchymal stromal cells (pBAT), immortalized BUC-derived cells (iBUC), myogenically induced BUC-derived cells (dBUC), primary satellite cells (pSAT), and differentiated satellite cells (dSAT) (Figure 1A-B; Supplementary Table S1). Primary and differentiated satellite cells were included as reference populations for myogenic differentiation.

The experimental design incorporated several biological dimensions, including tissue origin, primary versus immortalized state, culture architecture, passage group, and myogenic differentiation state (Figure 1A). Cells were represented in conventional two-dimensional plate culture, three-dimensional cellulose-based scaffold culture, and non-adherent spheroid culture (Figure 1A-B; Supplementary Table S1).

The primary cell models were characterized prior to their inclusion in the transcriptomic analysis. pBUC populations had been previously characterized for MSC-associated properties, including marker expression, proliferation, and multilineage differentiation capacity. pBAT cells were characterized in the present study for MSC-associated properties (Supplementary Figure S1A-C), and their growth and marker expression under spheroid culture conditions were also evaluated (Supplementary Figure S1D-E). Satellite cells were characterized under undifferentiated and myogenic conditions (Supplementary Figure S1F).

To assess passage and immortalization-associated transcriptional changes, pBUC and iBUC populations were compared at defined early and late stages of culture expansion, as detailed in the Methods and Supplementary Table S1. The complete RNA-seq dataset was subsequently analyzed jointly.

### Cell-cycle-related gene expression is a major source of variation between conditions and correlates with proliferation rates

Global transcriptional relationships revealed that culture architecture and cell identity were major sources of variation (Figure 2A). Biological replicates showed high within-condition similarity, with no apparent reduction in similarity among pBUC samples derived from three different cows compared with pBAT replicates obtained from a single cow. Distinct experimental states occupied separate regions of transcriptional space. Spheroid and immortalized populations occupied distinct transcriptional states, whereas plate-and scaffold-grown primary cells were more closely related, with scaffold-grown cells forming a distinguishable subgroup.

**Figure 2.**
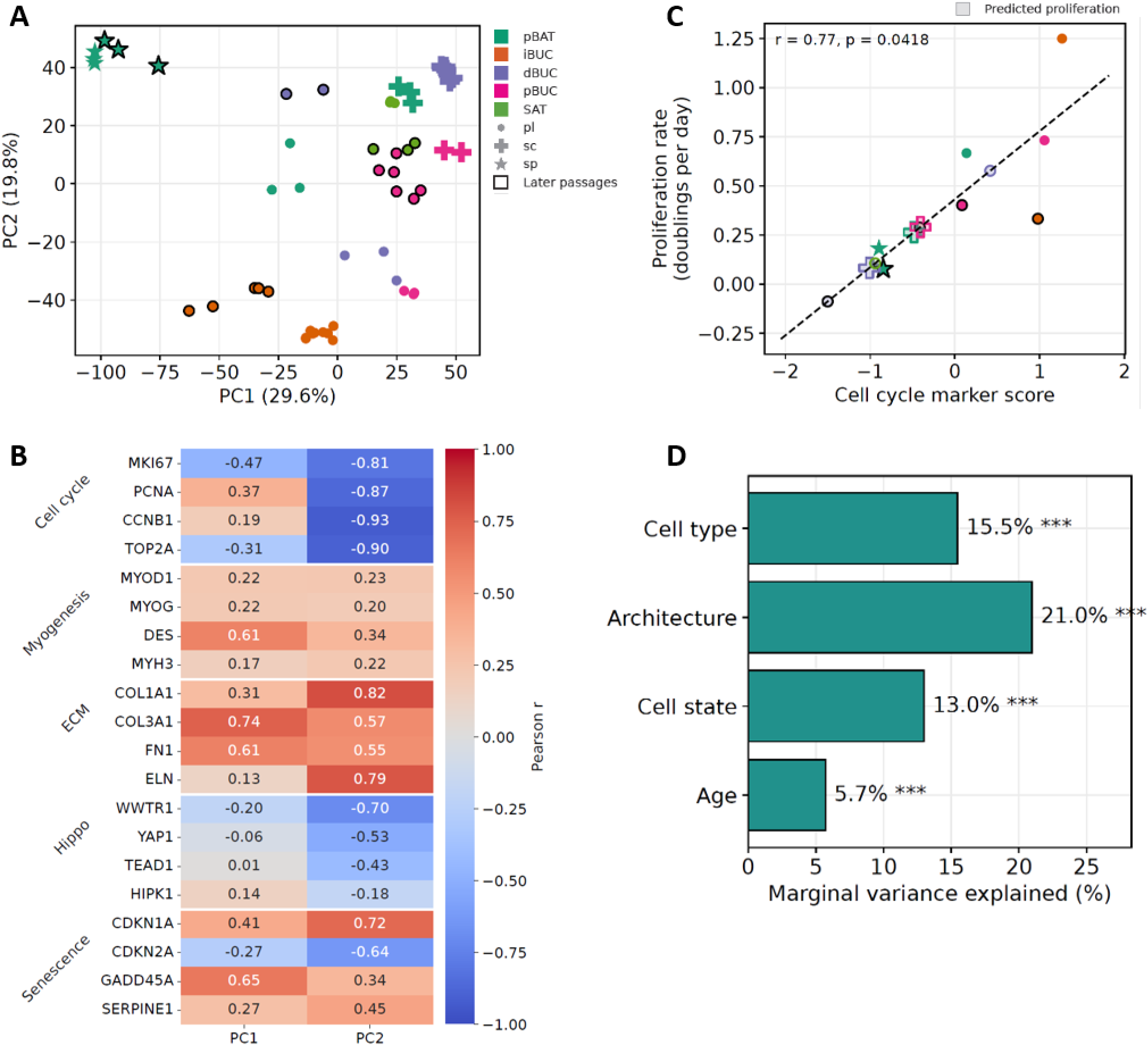
Global transcriptional variation and its relationship with cellular properties. **A.** PCA of DESeq2 variance-stabilized gene-expression counts. Each point represents one biological replicate; colors and shapes denote sample properties as indicated, and black outlines mark later passages. **B.** Pearson correlations between PC1 and PC2 sample scores and the expression of marker genes representing cell cycle, myogenesis, extracellular matrix, and senescence. **C.** Relationship between the mean standardized expression of the cell-cycle markers MKI67, PCNA, CCNB1, and TOP2A and measured proliferation rates. The dashed line shows the linear regression fitted to conditions with measured proliferation; shaded markers indicate proliferation rates estimated from this model. **D.** Marginal variance explained by culture architecture, cell type, cell state, and age, determined by PERMANOVA.

To identify the main drivers of transcriptional variation, we examined sample-level correlations between the first two principal components and the expression of selected marker genes. PC1 showed weak correlations with extracellular matrix (ECM)-associated genes, whereas PC2 was strongly correlated with several key cell-cycle markers, indicating that proliferative activity was a major contributor to transcriptional variation. PC2 was also correlated with genes associated with the Hippo signaling pathway, which integrates signals related to cell proliferation, cell–cell interactions, and mechanotransduction (Figure 2B). Consistent with the cell-cycle associations, experimentally measured proliferation rates were strongly correlated with enrichment of the cell-cycle biological process (GO:0007049; r = 0.90, p = 0.006; Supplementary Figure S2A).

We then calculated a cell-cycle score by averaging marker-gene expression for each sample and subsequently averaging across replicates within each condition. This score was significantly correlated with measured proliferation rates, with iBUCs showing the highest and BAT spheroids the lowest proliferation rates (Figure 2C). We fitted a linear model between the cell-cycle score and measured proliferation rate and used it to estimate proliferation rates for conditions in which proliferation was not measured, particularly the scaffold samples. These estimates suggested slow proliferation of scaffold-grown cells. Finally, we used PERMANOVA to attribute transcriptional variation to culture architecture, cell type, cell state, and age (Figure 2D). Together, these properties explained 73% of the global transcriptomic variation. Thus, culture architecture was associated with substantial transcriptional differences even after accounting for differences in cellular identity and state.

### Immortalization preserves early-passage-like features while opposing primary BUC aging

To determine how SV40LT-mediated immortalization alters the transcriptional state of BUCs, we compared early-and late-passage primary and immortalized populations. Transcriptome-wide relationships showed that iBUCs retained greater transcriptional similarity to early-passage pBUC_12pl than to late-passage pBUC_50pl, although this resemblance decreased with prolonged passage of the immortalized cells (Figure 3A, Supplementary Figure S3A). Pathway-level comparison further showed that late-passage iBUCs retained the proliferative and cell-cycle-related profile characteristic of early-passage BUCs and iBUCs, in contrast to the pathway profile associated with primary-cell aging (Figure 3B, Supplementary Figure S3B). Late-passage pBUCs instead showed broader similarities to scaffold-grown and differentiating populations, including increased representation of developmental and differentiation-related pathways (Figure 3C). Late-passage pBUCs also showed enrichment of inflammatory signaling, including NF-κB-associated pathways (Supplementary Figure S3C). Several inflammatory genes, including IL6, CXCL8, NFKB1, NFKBIA, and TLR2, also contributed to the aging-associated transcriptional response (Figure 3D).

**Figure 3.**
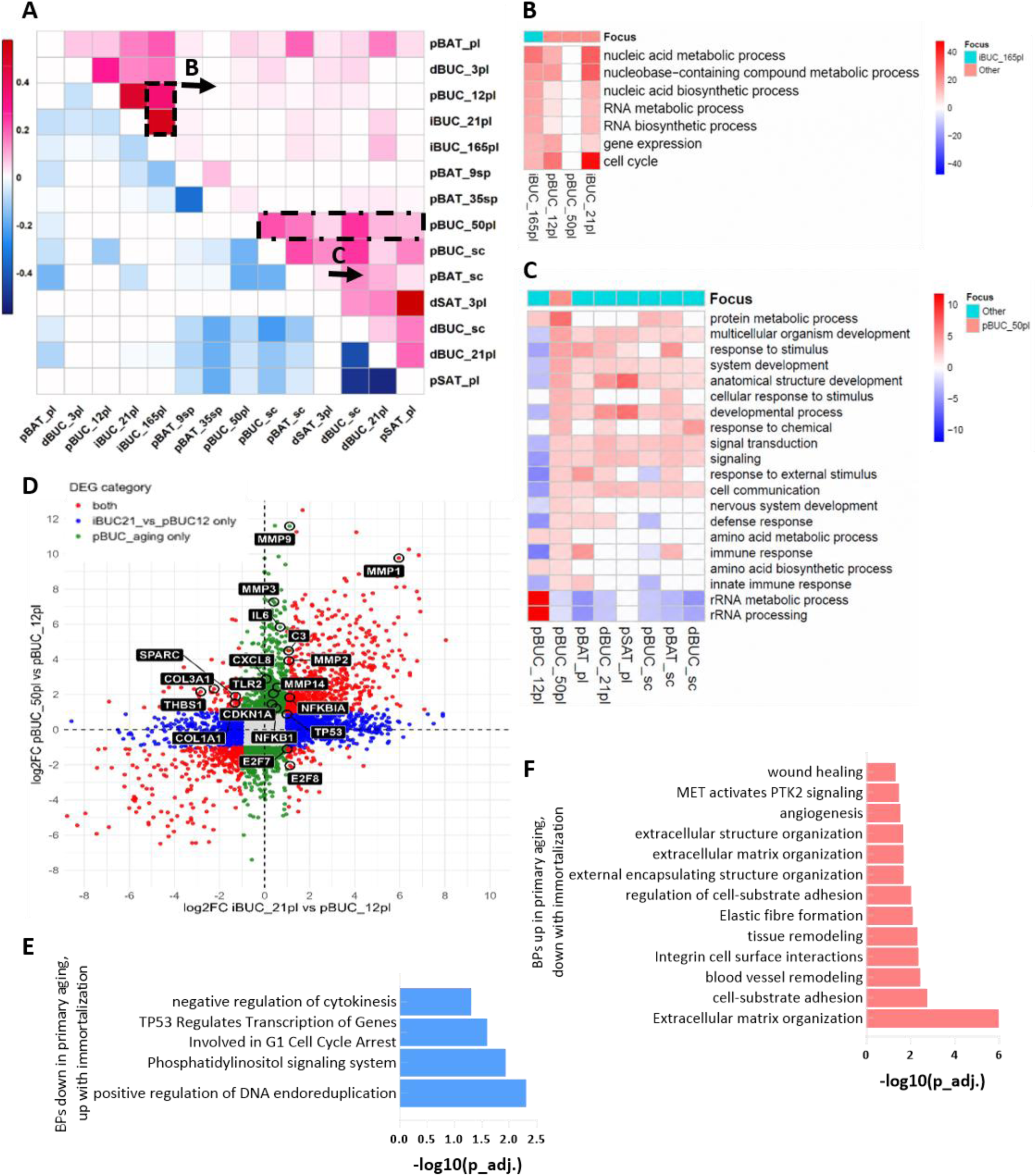
Immortalization preserves early-passage-like features while opposing primary BUC aging. **A.** Jaccard similarity of significantly enriched up-and downregulated Biological Processes identified by one-versus-all analysis. **B-C.** Enrichment profiles of selected shared Biological Processes across cell populations. Red indicates positive enrichment, blue negative enrichment, and white non-significant enrichment. Late-passage iBUC is the focus in B, and late-passage pBUC is the focus in C. **D.** Scatter plot comparing gene-level log2 fold changes associated with immortalization (iBUC_21pl vs pBUC_12pl) and primary BUC aging (pBUC_50pl vs pBUC_12pl). Red points indicate genes differentially expressed in both contrasts, blue points genes significant only in the immortalization contrast, and green points genes significant only in the primary-aging contrast. Selected genes associated with cell-cycle regulation, extracellular-matrix remodeling, and inflammatory signaling are labelled. Opposing quadrants highlight genes whose regulation differs in direction between immortalization and primary aging. **E.** Biological Processes up in primary aging / down with immortalization. Bars represent −log10(adjusted p-value). **F.** Biological Processes down in primary aging / up with immortalization. Bars represent −log10(adjusted p-value).

Despite these broader similarities, the four populations retained distinct transcriptional programs. Early-passage iBUCs were enriched for cell-cycle and genome-maintenance processes, whereas late-passage primary BUCs showed increased endoplasmic-reticulum stress, protein-quality-control, inflammatory-signaling, and extracellular-matrix-remodeling programs (Supplementary Table S2). Late-passage iBUCs retained mitotic activity but showed greater enrichment of metabolic, homeostatic, and stress-adaptive processes than iBUC_21pl, indicating progressive passage-dependent changes within the immortalized lineage.

Direct comparison of primary-cell aging and immortalization revealed both opposing and shared gene-level responses (Figure 3D), with pathway enrichment further highlighting opposing biological programs (Figure 3E–F). Cell-cycle-associated genes showed opposing regulation during immortalization and primary aging, including E2F7, E2F8, TP53, and CDKN1A (Figure 3D). Conversely, extracellular-matrix and tissue-remodeling genes including COL1A1, COL3A1, THBS1, and SPARC increased during primary aging but decreased following immortalization (Figure 3D). Consistent with these gene-level changes, enrichment analysis identified opposing regulation of extracellular-matrix organization, collagen-associated processes, cell-junction programs, and cell-cycle-related processes (Figure 3E-F, Supplementary Table S2). Immortalization was additionally associated with enrichment of cytokinesis-and DNA-endoreduplication-related processes (Figure 3F). Consistent with this transcriptional signature, immortalized cells displayed substantially increased chromosome numbers compared with non-immortalized cells (Supplementary Figure S3E-F). Despite these predominantly opposing responses, aging and immortalization also shared changes in selected remodeling-associated genes, including MMP1, MMP2, MMP3, MMP9, and C3 (Figure 3D). Together, these findings indicate that immortalization does not simply reverse primary-cell aging: it preserves selected early-passage-like features and opposes aging-associated loss of proliferative programs, while establishing a distinct proliferative and stress-adapted transcriptional state.

### Culture architecture shapes distinct transcriptional programs within pBAT

Because cell identity itself contributed strongly to global transcriptional variation (Figure 2D), we first examined culture architecture within a single cell type. pBAT cells were present in plate, scaffold, and spheroid cultures, which differed not only in spatial organization but also in adhesive environment, medium composition, and physical conditions (e.g., static vs. agitated culture for spheroids). We therefore treated these configurations as distinct culture systems rather than examining spatial geometry alone. The three culture formats occupied distinct transcriptional states, and within the pBAT subset culture architecture was associated with 80.4% of transcriptional variation (PERMANOVA, p = 0.001; Supplementary Figure S4A).Architecture-associated pathway analysis highlighted focal adhesion, adherens junction, Hippo signaling, mitophagy, ubiquitin-mediated proteolysis, and signaling pathways regulating stem-cell pluripotency (Supplementary Figure S4B).

Architecture-associated DEGs resolved into distinct transcriptional programs across the three culture conditions (Figure 4B). These included extracellular-matrix and collagen organization, cell-cycle regulation, endocytosis and autophagy, TGF-β associated signaling, and lipid and lipoprotein metabolism. Scaffold-associated Biological Processes were particularly enriched for extracellular-structure organization and tissue remodeling, collagen fibril organization, response to endoplasmic-reticulum stress, regulation of cell motility, and developmental programs (Figure 4C). In contrast, spheroids showed coordinated downregulation of transcription factors linked to developmental and differentiation-related programs (Supplementary Figure S4C). Despite their distinct transcriptional states, scaffold and spheroid cultures shared several changes relative to plate culture, including increased extracellular-matrix and collagen-associated and integrin-related programs and mTOR signaling, together with reduced cell-cycle-associated enrichment (Figure 4D).

**Figure 4.**
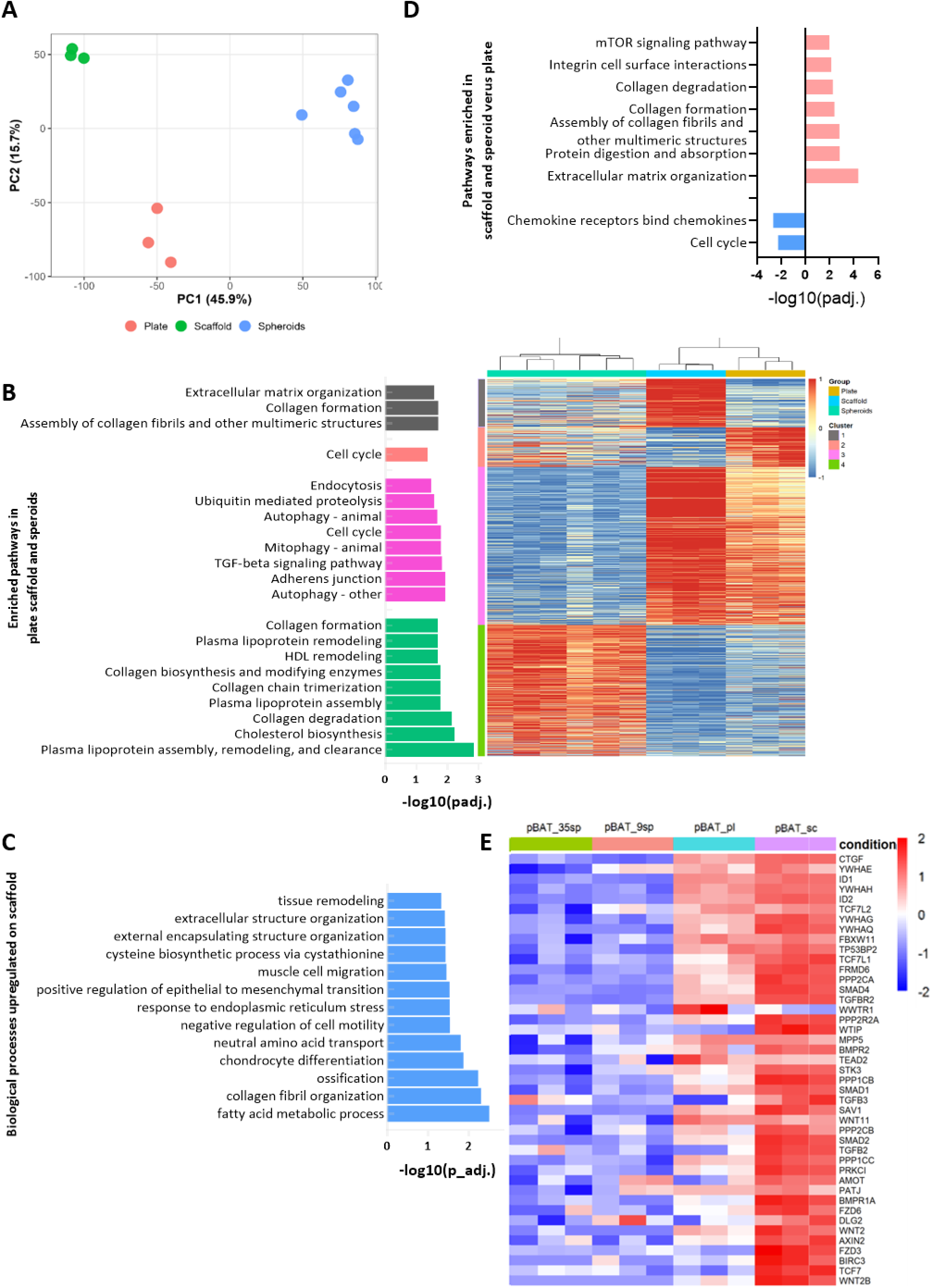
Culture architecture shapes distinct transcriptional programs within pBAT. **A.** Global transcriptional relationships among pBAT cells cultured on plate, scaffold, or as spheroids, showing separation of the three culture architectures. **B.** Architecture-associated differentially expressed genes organized into transcriptional programs, with representative enriched pathways shown for the corresponding expression patterns. Enrichment is represented as −log10(adjusted p-value). **C.** Biological Processes enriched among scaffold-associated genes. Bars represent −log10(adjusted p-value). **D.** Pathways commonly altered in scaffold and spheroid cultures relative to plate culture. Positive values indicate enrichment among upregulated genes and negative values enrichment among downregulated genes. Only terms with |signed −log10(adjusted p-value) | > 2 are shown. E. Expression of selected Hippo-signaling and mechanotransduction-related genes across pBAT culture architectures. Expression values are shown as row-scaled Z-scores.

Direct comparison of pBAT scaffold-versus-plate and spheroid-versus-plate differential-expression responses further emphasized their divergence. The two contrasts were slightly negatively correlated (r = −0.15, p = 1 × 10⁻⁶⁷; Supplementary Figure S4D), consistent with substantial differences in the direction of transcriptional responses between the two 3D environments. Extracellular-matrix-associated programs increased in both conditions, whereas endocytosis showed opposite behavior, increasing in scaffold culture but decreasing in spheroids (Figure 4B, Supplementary Figure S4D).

Consistent with the enrichment of adhesion-and mechanotransduction-associated pathways, Hippo-and mechanotransduction-related genes showed a clear architecture-dependent expression pattern (Figure 4E). Scaffold and spheroid cultures showed non-identical expression profiles across this gene set, further supporting the conclusion that the transcriptional response depended on the specific culture environment rather than representing a uniform response to three-dimensional culture.

Within spheroid cultures, prolonged culture was also associated with a distinct transcriptional shift. In the 35-day versus 9-day comparison, enrichment analysis identified increased representation of lysosome-, N-glycan-biosynthesis-, and protein-metabolism-related pathways, whereas genes associated with DNA metabolism, DNA replication, nucleosome organization, and chromatin organization were reduced (Supplementary Figure S4E). The latter included chromatin-regulatory genes such as SMARCA2, SMARCC1, SMARCD1, KAT7, KMT2A, PHF8, and ARID4B (Supplementary Table S2). Together, these changes are consistent with reduced proliferative and chromatin-associated activity during prolonged spheroid culture, accompanied by increased protein-processing and intracellular-remodeling functions.

### Scaffold culture induces recurrent but cell-state-dependent transcriptional responses

The broader pathway comparison revealed that multiple scaffold-grown populations shared developmental and extracellular-matrix-associated programs, with a similar pattern also observed in late-passage pBUCs (Figure 3C). We therefore asked whether these responses were conserved across different cell states. Having identified scaffold-associated programs within the controlled pBAT comparison, we compared scaffold-grown pBUC, pBAT, and dBUC (cells after 21 days of myogenic differentiation) populations with their corresponding plate-grown controls. Differential-expression responses were significantly correlated across the three scaffold-versus-plate contrasts, with the strongest similarity between pBAT and pBUC and weaker correlations in comparisons involving dBUC (Figure 5A–B). Shared scaffold-associated genes included extracellular-matrix and adhesion genes such as ELN, COL16A1, and CDH11, together with FZD1 and GJA1, indicating recurrent changes in matrix organization, adhesion, and intercellular signaling.

**Figure 5.**
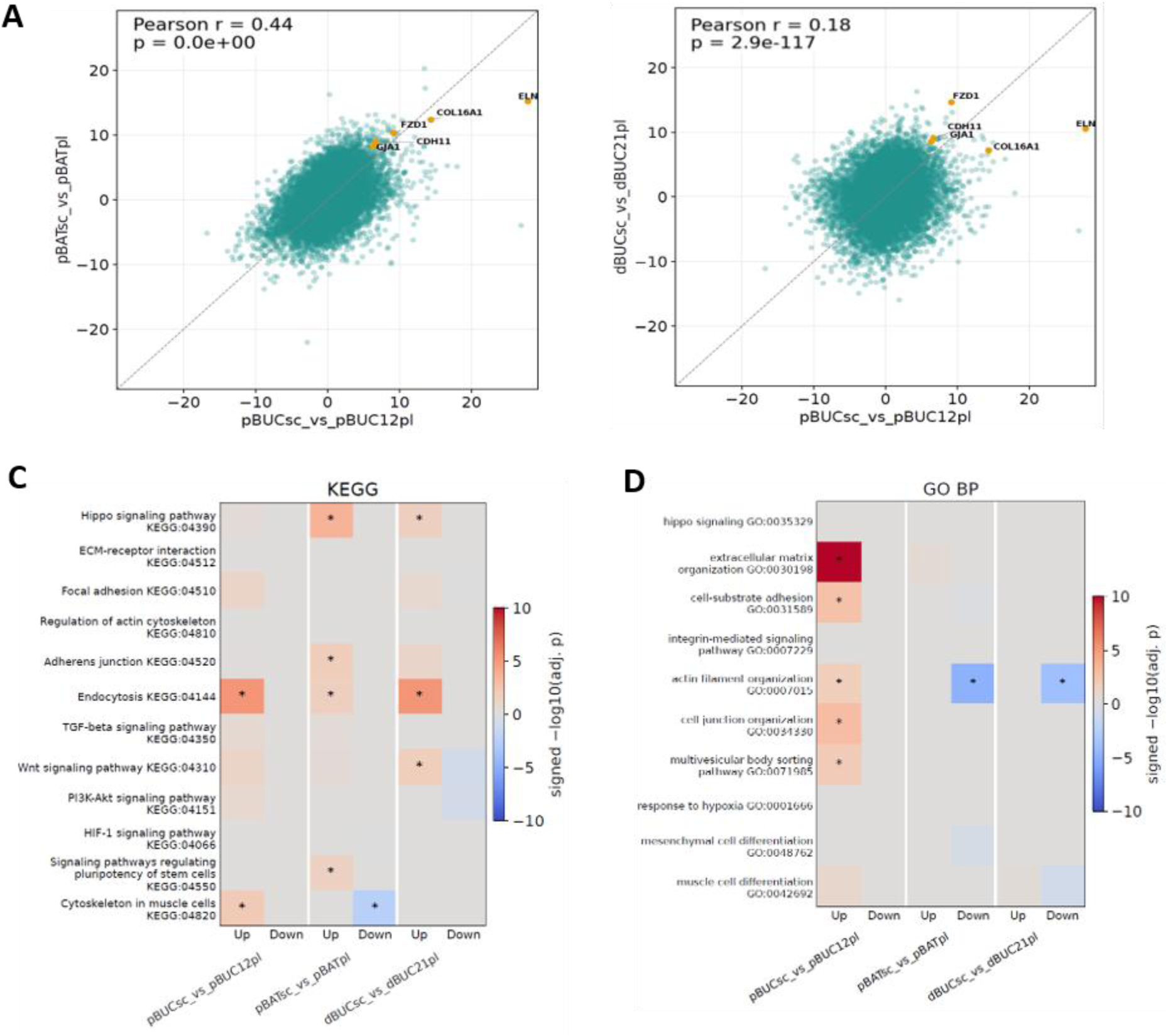
Transcriptional responses to scaffold versus plate culture. A-B. Pairwise comparison of DESeq2 Wald statistics for scaffold-versus-plate contrasts across cell types, with Pearson correlation coefficients. **C-D.** Signed enrichment scores for a manually selected, non-redundant set of KEGG pathways and GO biological process terms related to cell structure, adhesion, mechanotransduction, and signaling across the three scaffold-versus-plate comparisons. Scores represent −sign (NES) × log10(adjusted p-value); asterisks indicate adjusted p < 0.05.

Functional enrichment revealed both common and cell-state-specific responses (Figure 5C). Several signaling pathways, including Wnt, TGF-β, PI3K–Akt, and HIF-1, showed scaffold-associated changes in individual cell states but were not consistently significant across all three comparisons. Hippo signaling showed positive enrichment in pBAT and dBUC scaffold cultures but was not consistently significant across all three comparisons, indicating a cell-state-dependent transcriptional response within this pathway. In contrast, endocytosis was the only selected pathway significantly enriched across all three scaffold-versus-plate comparisons (Figure 5C).

Pathway-level visualization showed recurrent regulation of genes involved in endosomal trafficking, recycling, vesicle formation, and endosomal sorting, including RAB4, RAB5, RAB11, RAB22, DAB2, VPS26, VPS35, and VPS37, whereas AMPH and endophilin-associated components were reduced (Supplementary Figure S5). The consistency of this response across pBUC, pBAT, and dBUC suggests that altered membrane and receptor trafficking is a recurrent feature of scaffold culture, potentially accompanying remodeling of cell–matrix and cell–cell interactions.

Taken together, the pBAT comparison demonstrates that culture architecture can strongly reshape the transcriptional state within a single cell type, while the cross-cell-state analysis identifies a subset of responses that recur in scaffold culture across distinct cellular contexts. The data therefore distinguish architecture-specific responses, including the marked differences between scaffold and spheroid cultures, from recurrent scaffold-associated programs, particularly endocytosis, extracellular-matrix remodeling, adhesion, developmental and differentiation programs, and selected mechanotransduction pathways.

### Myogenic programs are partially shared between BUC-derived and satellite-cell differentiation and are supported by scaffold culture

To assess how culture architecture influences myogenic differentiation-related transcriptional programs, we compared BUC cells cultured on conventional plates and cellulose-based scaffolds under undifferentiated and myogenically induced conditions. Primary bovine satellite cells, which originate from skeletal muscle and represent endogenous myogenic progenitors, were included as a canonical myogenic reference. This design allowed us to interpret BUC differentiation programs in the context of established skeletal-muscle myogenesis while directly evaluating the contribution of culture architecture.

Global transcriptional relationships remained strongly influenced by cell identity, with BUC-derived and satellite-cell-derived populations occupying distinct transcriptional states (Figure 6A). Consistent with the PCA structure, PERMANOVA showed significant associations of cell source, myogenesis stage, and culture architecture with global transcriptomic variation, explaining 30%, 21%, and 16% of the variation, respectively (all *p* = 0.001). Satellite cells also showed stronger expression of selected myogenic markers and a distinct myogenic transcription-factor profile compared with BUC-derived cells (Supplementary Figure S6A–B). Nevertheless, myogenic induction shifted both cell types away from their respective undifferentiated states, indicating differentiation-associated transcriptional remodeling in both systems (Figure 6A).

**Figure 6.**
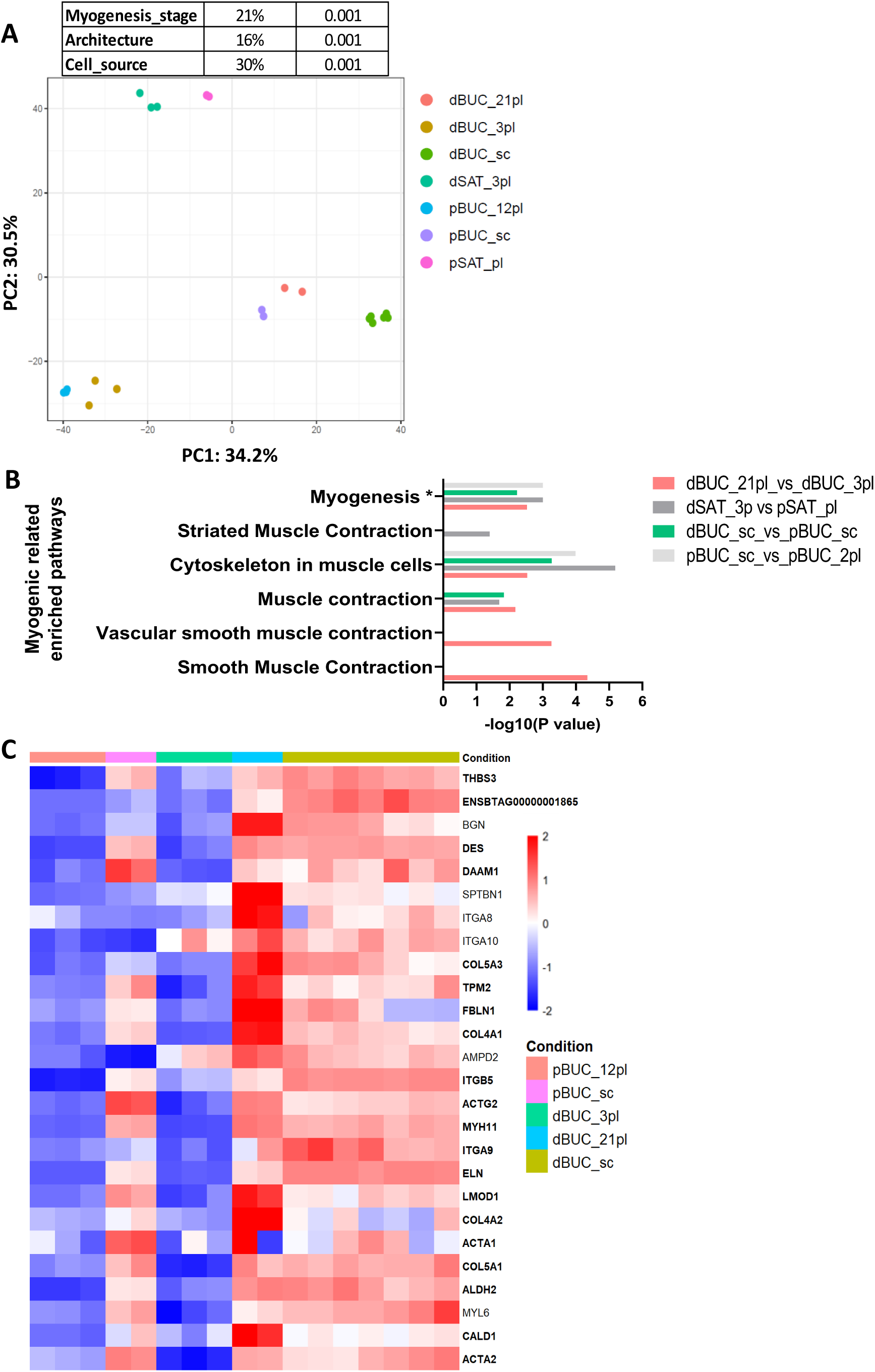
BUC myogenic induction partially overlaps with satellite cell differentiation, and scaffold culture is associated with myogenesis-related transcriptional signatures. **A.** Global transcriptional relationships among plate-cultured undifferentiated and myogenically induced BUCs and satellite-cell-derived cultures. **B.** Shared and distinct myogenesis-related pathway enrichment during BUC-derived and satellite-cell differentiation. Pathway enrichment was performed using g-profiler on differentially expressed genes, except for Hallmark Myogenesis (*). Bars represent −log10(adjusted p-value) and −log10(nominal p-value) for GSEA. **C.** Expression of genes associated with Cytoskeleton in muscle cells (KEGG:04820) and Smooth Muscle Contraction (REAC:R-BTA-445355) showing differentiation-associated changes in BUCs cultured on plates and/or scaffolds. Genes shown in bold are DEGs between pBUC_sc and pBUC_pl.

Comparison of differentiation-associated programs revealed both shared and cell-type-specific responses. In both BUCs and satellite cells, differentiation was associated with reduced cell-cycle-related programs and increased muscle-, cytoskeletal-, and extracellular-matrix-associated processes (Supplementary Figure S6C–E). Several muscle-related pathways were enriched in both differentiated populations, supporting partial convergence toward a myogenic transcriptional state (Figure 6B). Consistent with this, GSEA of the ranked transcriptomic data revealed positive enrichment of the Hallmark Myogenesis gene set, further supporting the presence of a broader myogenesis-associated transcriptional signature (Figure 6B). However, the nature of these responses differed between the two cell sources. Differentiated satellite cells showed stronger enrichment of striated-muscle and skeletal-muscle-contraction programs, whereas differentiated BUCs showed broader enrichment of cytoskeletal, extracellular-matrix-remodeling, adhesion, and smooth-muscle-like contractile programs. Thus, BUC-derived myogenic induction reproduced part of the transcriptional program observed during satellite-cell differentiation while retaining a distinct BUC-associated transcriptional identity, consistent with the strong contribution of cell source to the global transcriptional differences between the two populations.

We next examined whether scaffold culture was associated with myogenic-related transcriptional changes in BUCs independently of myogenic induction. In the global transcriptional space, undifferentiated scaffold-grown BUCs were displaced from undifferentiated plate-grown BUCs and positioned closer to differentiated BUC populations, consistent with an architecture-associated shift toward a differentiation-related transcriptional state (Figure 6A). Notably, GSEA revealed positive enrichment of the Hallmark Myogenesis gene set in undifferentiated scaffold-grown BUCs relative to plate-grown BUCs, indicating that scaffold culture alone was associated with a broader myogenesis-related transcriptional signature (Figure 6B). To examine this relationship at the gene level, we focused on genes belonging to the “cytoskeleton in muscle cells” and “smooth muscle contraction” pathways (Figure 6B). Many of these genes were differentially expressed between undifferentiated BUCs grown on scaffold and those grown on conventional plates, including genes with expression patterns similar to those observed following myogenic induction (Figure 6C). These results indicate that scaffold culture alters muscle-and cytoskeleton-associated transcriptional programs even in the absence of myogenic differentiation medium.

Together, these findings indicate that BUC-derived myogenic induction shares core muscle-and cytoskeleton-associated transcriptional programs with satellite-cell differentiation, despite remaining distinct from the more specialized skeletal-muscle program of satellite cells. In parallel, scaffold culture alters the basal BUC transcriptional state and is associated with positive enrichment of a broader myogenesis-related transcriptional program, supporting a role for culture architecture in promoting myogenesis-associated transcriptional features in BUCs.

## Discussion

Cell culture is often optimized one variable at a time: cell source, number of passages, growth format, or differentiation protocol. Yet these choices are not independent, and cell selection and culture-system design are closely linked challenges in cultivated meat development. Here, we integrated transcriptomic data from bovine stem and progenitor cells from different tissues, at different passages and differentiation states, grown on plates, in spheroids, or on cellulose scaffolds. Most of these cells had already been characterized in our previous studies, allowing us to examine their responses within a common analytical framework. Our findings distinguish shared features of cellular state from adaptations associated with particular culture environments. The most prominent shared changes involved proliferation-associated programs, whereas differences in culture architecture and cellular identity were reflected in pathways related to ECM organization, membrane trafficking, metabolism, and differentiation.

The comparison of plate, scaffold, and spheroid cultures revealed considerable differences between the two 3D systems (Figure 4). Cells attached to cellulose scaffolds showed changes in ECM organization, adhesion, and membrane trafficking. Spheroid-grown BAT cells also showed ECM-associated changes but had a stronger reduction in cell-cycle and chromatin-related programs, together with changes in lysosomal and protein-processing pathways. Interestingly, endocytosis-associated pathways were regulated in opposite directions in scaffold and spheroid cultures. These differences should be interpreted in the context of the complete culture configurations. Throughout this study, culture architecture encompasses not only spatial organization but also the attachment environment, medium formulation, and physical conditions used to establish and maintain each format. Scaffold cultures provide a solid substrate for attachment, whereas spheroid cultures combine non-adherent conditions with a supporting medium and agitation. Our comparisons therefore address the transcriptional states associated with these integrated culture systems, rather than the isolated effect of three-dimensional geometry. This system-level perspective is relevant to cultivated meat development, where cells must function within a complete set of culture conditions. A complementary question for future studies is how individual components, and their interactions, contribute to the distinct responses observed here.

Proliferation is an important factor when comparing cell populations with different passage histories and culture conditions. As expected, cell-cycle gene expression was strongly correlated with measured proliferation rates (Figure 2; Supplementary Figure S2). This correlation allowed us to estimate growth rates for conditions in which proliferation was not directly measured, although these estimates require experimental validation. More importantly, reduced proliferation-associated transcription was observed under several otherwise different conditions, including prolonged passaging of primary BUCs and cultivation on scaffolds or as spheroids. However, the other transcriptional changes accompanying reduced proliferation differed considerably. This was particularly evident for developmental programs, which were increased in late-passage primary BUCs and scaffold cultures but decreased in spheroids. Late-passage primary BUCs showed increased inflammatory and stress-related programs, whereas spheroid cultures displayed pronounced changes in chromatin organization and metabolism. Thus, conditions with similarly reduced proliferation could engage distinct, and in some cases opposing, transcriptional programs. This suggests that proliferation-related signatures may explain part of the similarity between transcriptional profiles without necessarily reflecting similar cellular states. Accounting for these differences could be particularly useful when comparing datasets generated under different culture conditions. In addition, passage number alone may be an insufficient measure of culture history, as cells with different proliferation rates may have undergone different numbers of divisions during the same period in culture.

The comparison between primary and immortalized BUCs further illustrates the importance of considering cellular properties beyond proliferation (Figure 3). Immortalized cells maintained high proliferative activity during prolonged culture and retained several transcriptional features characteristic of early-passage primary cells. However, they also displayed changes in metabolic and stress-related pathways, together with enrichment of genes associated with cytokinesis and DNA endoreduplication. These findings are consistent with the increased chromosome numbers observed in the immortalized population, although the mechanisms underlying these changes remain unclear. Interestingly, certain ECM-associated changes were shared between late-passage primary and immortalized BUCs, suggesting that prolonged culture may produce some similar responses despite their different proliferative capacities. Immortalization can overcome the limited expansion potential of primary cells, which is particularly advantageous for applications requiring large quantities of cells. However, our results emphasize the need to characterize immortalized lines beyond their growth properties, including genomic stability and maintenance of differentiation potential. It would also be interesting to determine whether the observed changes are specific to the SV40LT immortalization strategy used here or represent more general adaptations associated with long-term expansion.

We next examined whether cells of different origins and differentiation states show common responses to cellulose scaffolds. The transcriptional response was more similar between the two undifferentiated MSC populations than between either of these populations and myogenically induced BUCs (Figure 5). Nevertheless, several pathways were recurrently affected, particularly those associated with endocytosis and intracellular trafficking. Endocytosis was among the most consistently enriched pathways across the three scaffold comparisons, suggesting that membrane trafficking may have an important role in cellular adaptation to scaffold culture. Endocytosis and membrane recycling regulate the localization and availability of cell-surface receptors, including integrins, and are involved in the continuous remodeling of cell–matrix interactions (Caswell et al., 2009; Moreno-Layseca et al., 2019). We suggest that the observed changes may be related to the reorganization of cell attachment and signaling when cells are grown on the porous cellulose structure. This possibility is supported by accompanying changes in ECM organization and adhesion-related pathways. However, pathway enrichment does not establish increased endocytic activity, and we cannot determine which receptors or trafficking mechanisms are involved. Future experiments examining integrin localization, internalization, and recycling could help clarify the role of these pathways in scaffold adaptation. Testing additional scaffold materials would also determine whether this response is specific to cellulose or represents a more general feature of cells grown on three-dimensional substrates.

The myogenic comparisons provided additional insight into the influence of cellular identity on culture responses (Figure 6). Both myogenically induced BUCs and differentiated satellite cells showed increased expression of muscle-associated genes, although the differentiated satellite cells exhibited stronger skeletal-muscle-specific signatures. Induced BUCs retained a broader transcriptional profile, including cytoskeletal, adhesion, and smooth-muscle-like contractile programs. These findings are consistent with our previous characterization of BUC-derived cells and suggest that the induction protocol activates selected myogenic programs without fully reproducing the transcriptional state of differentiated satellite cells. Interestingly, undifferentiated BUCs grown on cellulose scaffolds also showed enrichment of myogenesis-associated gene sets, even without myogenic induction medium. This observation is consistent with our previous findings that cellulose scaffolds can support myogenic differentiation of bovine MSCs (Gershkoviz et al., 2026). However, many genes included in myogenesis-related pathways are also involved in general cytoskeletal organization and cell adhesion. Additional experiments are therefore required to determine whether scaffold culture promotes skeletal-muscle differentiation or mainly induces changes associated with cytoskeletal remodeling. In particular, muscle-specific protein expression, myotube formation, and functional maturation should be examined. It should also be considered that BUCs and satellite cells originate from different tissues and were subjected to different differentiation protocols, limiting direct comparisons of their myogenic competence.

Several limitations should be considered when interpreting these findings. First, the study compared integrated culture configurations rather than isolating the contributions of substrate attachment, medium composition, and physical forces. The observed responses therefore relate to the complete culture systems and cannot be attributed to individual components. Second, the dataset combined cell populations from different projects within a non-factorial experimental design. Although RNA extraction and sequencing were performed together, differences in donor background, culture duration, passage history, and project-specific handling may have contributed to the observed variation. Not all combinations of cell source, differentiation state, passage, and culture architecture were represented, limiting our ability to disentangle their respective contributions. In particular, BAT cells originated from a single donor, and spheroid culture was examined only in this population, restricting the generalizability of these findings. Third, bulk RNA-seq cannot distinguish changes in gene expression within cells from shifts in the relative abundance of different cell states, a distinction that may be particularly relevant in spheroid cultures. Finally, interpretations concerning signaling, trafficking, and differentiation are based primarily on transcriptional profiles and pathway enrichment rather than direct functional measurements. Studies incorporating additional donors, matched comparisons within each culture configuration, and targeted perturbations of individual culture components will be needed to test the proposed mechanisms and establish how broadly these responses extend to other cell populations and culture systems.

Our findings have implications for the development of cell culture systems for tissue engineering, regenerative medicine, and cellular agriculture. These applications often require prolonged cell expansion followed by changes in culture conditions to promote differentiation or tissue formation. In cultivated meat production, for example, cells may be expanded in suspension before being transferred to scaffolds for maturation. Our results suggest that the performance of a given cell population should be evaluated under the different conditions it is expected to encounter, as its proliferative capacity and differentiation potential under conventional plate culture may not predict its response to a different architecture. The recurrent changes in membrane trafficking and adhesion-related pathways may also provide a direction for understanding and improving cell–scaffold interactions, although their functional contribution remains to be established. Overall, this study distinguishes recurrent transcriptional adaptations from responses specific to bovine cell source, expansion history, and culture system. By identifying both shared scaffold-associated trafficking signatures and divergent responses to different three-dimensional environments, it provides a framework for evaluating cell–culture combinations for cultivated meat and prioritizing mechanisms for functional validation.

## Methods

### Cell culture

Bovine umbilical cord-derived mesenchymal stromal cells

Bovine umbilical cord-derived mesenchymal stromal cells (BUCs) were isolated, cultured, and characterized according to generally accepted MSC criteria (Dominici et al., 2006; Uccelli et al., 2008), as previously reported (Ribarski-Chorev et al., 2023; Nir et al., 2022; Shimoni et al., 2020). Briefly, umbilical cords from Holstein dairy cows were obtained from abattoirs located in northern Israel. In the laboratory, umbilical cords were digested and plated as previously described by Toupadakis et al. (2010). Cells were expanded in low-glucose Dulbecco’s Modified Eagle’s Medium (DMEM; Gibco, 31885-023, or Biological Industries, 04-001-1A) supplemented with 10% fetal bovine serum (FBS), 1% penicillin-streptomycin, and 1% L-glutamine. Cells were cryopreserved at different passages in FBS containing 10% DMSO (Sigma-Aldrich, W387520). Medium was changed every 2–3 days, and cells were maintained at 37°C in a humidified atmosphere containing 5% CO₂ unless otherwise stated.

### Bovine adipose tissue-derived mesenchymal stromal cells

Bovine adipose tissue-derived mesenchymal stromal cells (BATs) were isolated from adipose tissue using the protocol described by Oyarzo et al. (2021), with minor modifications. Briefly, tissue was immersed in 70% ethanol for 30 s and subsequently washed in Dulbecco’s phosphate-buffered saline (DPBS) containing 3% penicillin-streptomycin, 1% gentamicin, and 1% amphotericin. Tissue was minced into small pieces and digested with 0.5% type I collagenase (Gibco) at 37°C for 1 h with shaking. Collagenase activity was terminated by addition of culture medium, and the cell suspension was filtered through a 70-μm cell strainer. The filtrate was washed once with DMEM, plated in BUC growth medium supplemented with 10 ng/mL bFGF, and maintained at 37°C under 5% CO₂, with medium changes every 2–3 days.

For spheroid generation, six-well plates were coated with Biofloat™ (Facellitate) to prevent cell adhesion. Plates were incubated with Biofloat for 3 min at room temperature, excess solution was removed, and plates were allowed to dry for 1 h in a biological safety cabinet. Cells were subsequently cultured in NutriStem medium (Sartorius) and maintained on an orbital shaker inside the incubator.

### Bovine satellite cells

Bovine satellite cells were isolated from hind-limb skeletal muscle obtained from Holstein dairy cows at abattoirs in Israel. Tissue was placed in DPBS supplemented with 3% penicillin-streptomycin and transported to the laboratory within 3 h. Tissue processing was performed according to Ben-Arye et al. (2020).

Briefly, tissue was immersed in 70% ethanol for 30 s and subsequently washed in DPBS containing 5% penicillin-streptomycin, 1% gentamicin, and 1% amphotericin. Blood vessels, adipose tissue, and connective tissue were removed, and the muscle was minced into small pieces. Tissue was digested for 1.5 h at 37°C in a solution containing 60 mg collagenase type I in 20 mL DPBS supplemented with 1% penicillin-streptomycin, with vigorous shaking every 10 min.

Following digestion, the suspension was centrifuged at 200 × g for 10 min. The supernatant was transferred to a separate tube, while the remaining material was treated with pre-warmed 0.25% trypsin-EDTA at 37°C for 20 min. The supernatant fraction was centrifuged at 300 × g for 7 min, and the cell pellet was resuspended in AmnioMAX-II medium (Thermo Fisher Scientific, 11269016). Following trypsinization, the second fraction was centrifuged, resuspended in AmnioMAX-II medium, and combined with the first cell fraction. The pooled cells were centrifuged at 500 × g for 10 min and resuspended in AmnioMAX-II medium.

The cell suspension was passed through a 40-μm strainer into 100-mm culture dishes and incubated overnight at 37°C under 5% CO₂. Non-adherent cells, including satellite cells, were subsequently collected and transferred to 0.1% gelatin-coated tissue-culture plates containing fresh AmnioMAX-II medium.

### Two-dimensional and three-dimensional culture

BATs were maintained at 37°C in a humidified atmosphere containing 5% CO₂, with medium changes every 2–3 days. For conventional plate culture, pBAT cells were maintained in low-glucose DMEM supplemented with 10% FBS, 1% penicillin-streptomycin, and 10 ng/mL bFGF. For spheroid culture, cells were seeded into Biofloat-coated six-well plates as described above and maintained in NutriStem medium on an orbital shaker. Scaffold-based culture is described below.

### Cell characterization

The primary bovine mesenchymal cell populations used in this study were characterized for MSC-associated properties. Three primary BUC lines had previously been examined and validated (Nir et al., 2022; Shimoni et al., 2020), whereas the primary adipose-derived MSC population (pBAT) was characterized in the present study (Supplementary Figure S1).

Multilineage differentiation potential was assessed toward osteogenic, adipogenic, and chondrogenic lineages.

For osteogenic differentiation, cells were cultured in low-glucose DMEM containing 10% FCS, 10⁻⁷ M dexamethasone, 100 μM ascorbic acid-2-phosphate, and 10 mM β-glycerophosphate. Medium was replaced every 3 days, and differentiation was assessed after 10 days by Alizarin Red S staining.

For adipogenic differentiation, cells were cultured in low-glucose DMEM supplemented with 10% FBS, 10⁻⁶ M dexamethasone, 0.5 mM 3-isobutyl-1-methylxanthine, 0.2 mM indomethacin, and 10 μg/mL recombinant human insulin. Adipogenic differentiation was assessed using 0.3% Oil Red O in isopropanol or BODIPY 493/503 (Invitrogen), with nuclei counterstained using 5 μg/mL DAPI.

For chondrogenic differentiation, cells were cultured for 3 weeks in DMEM/F-12 supplemented with 5% FBS, 100 IU/mL penicillin, 0.1 mg/mL streptomycin, 2 mM L-glutamine, 1% insulin-transferrin-selenium (ITS-G; Gibco), and 10 ng/mL TGF-β1 (PeproTech). Chondrogenic differentiation was assessed by staining with 0.6% Alcian Blue.

Following staining, samples were evaluated microscopically using an EVOS FL Auto imaging system (Thermo Fisher Scientific). Negative controls consisted of stained cells maintained without differentiation medium. Unless otherwise specified, characterization experiments were performed using cells at passages 2–4 with at least three biological replicates.

### Characterization of BAT spheroids

BAT cells cultured under 2D and spheroid (3D) conditions were characterized by population doubling time and mesenchymal stromal cell marker expression. Population doubling time was monitored over time, and expression of positive and negative mesenchymal marker genes was assessed by RT–qPCR and normalized to the reference gene (Supplementary Figure S1D–E).

### Characterization of satellite cells

Satellite cells were evaluated under undifferentiated and day-3 myogenic differentiation conditions by bright-field microscopy and immunofluorescence staining. Cells were stained for myogenin (MyoG) or myosin heavy chain (MHC), together with phalloidin-labelled actin and DAPI nuclear counterstaining (Supplementary Figure S1F).

### SV40-immortalized BUCs

The immortalized bovine MSC line was generated as described by Tawil et al. (2026) by transient transfection of primary MSCs with a plasmid encoding the SV40 large T antigen, kindly provided by Prof. Sara Selig. Transfections were performed using TransIT-LT1 transfection reagent (Mirus Bio, MIR 2300) according to the manufacturer’s instructions. Colonies were expanded and subsequently assessed for proliferation, morphology, and genomic stability.

### Myogenic differentiation

Myogenic differentiation of BUCs

MSCs were seeded at 25,000 cells per well in six-well plates (2,600 cells/cm²) and maintained for 24 h in growth medium before initiation of differentiation (Gershkoviz et al., 2026). Myogenic differentiation was induced by replacing growth medium with DMEM/F-12 supplemented with 2% FBS, 1% ITS, 1% BSA, 10⁻⁸ M dexamethasone, 10⁻⁴ M ascorbic acid-2-phosphate (Sigma-Aldrich, A8960), 10 ng/mL vascular endothelial growth factor (PeproTech, 100-20), 10 ng/mL insulin-like growth factor (PeproTech, 100-11), 10 ng/mL fibroblast growth factor (PeproTech, AF-250-62), 1% penicillin-streptomycin, and 1% L-glutamine, based on Alviano et al. (2007). Medium was changed every other day for 21 days.

### Myogenic differentiation of satellite cells

Satellite cells were maintained in low-glucose DMEM supplemented with 20% FBS, 1% L-glutamine, 1% penicillin-streptomycin, and 1 ng/mL recombinant bovine FGF2 (PeproTech, AF-450-62), as described by Stout et al. (2023). Before induction of differentiation, cells were detached using 0.25% trypsin and seeded at 150,000 cells per well in 0.1% gelatin-coated six-well plates. Upon reaching confluence, medium was replaced with low-glucose DMEM supplemented with 2% horse serum, 1% penicillin-streptomycin, 1% L-glutamine, 100 ng/mL recombinant human IGF2, and 62 ng/mL recombinant human EGF, as described by Ianovici et al. (2022). Cells were differentiated for 3 days, and samples were collected on days 1 and 3 for RNA extraction.

### Cellulose scaffolds

Scaffold preparation, cell seeding and culture, and myogenic differentiation on cellulose-based scaffolds were performed as described by Gershkoviz et al. (2026).

### RNA sequencing

The RNA-seq samples analyzed in this study were generated across several related experimental projects. Cell pellets from all experimental groups were stored at −80°C until processing. RNA extraction and quality assessment were subsequently performed for all samples together in a single batch by the same researcher using standardized protocols, and all RNA samples were submitted together for library preparation and sequencing at the Nancy and Stephen Grand Israel National Center for Personalized Medicine, Weizmann Institute of Science.

Subsets of these samples were previously deposited in the Gene Expression Omnibus under accession numbers GSE276658 and GSE327833. These samples were analyzed here together with additional samples deposited under GSE329893, comprising the complete RNA-seq dataset analyzed in this study. A detailed description of the RNA-seq sample groups, including cell type, tissue origin, primary or immortalized status, culture architecture, passage group, and differentiation status, is provided in Supplementary Table S1.

For the analysis of passage-associated transcriptional changes, primary and immortalized BUC populations were divided into early-and late-passage groups based on their culture history. Early-passage pBUCs included passages 3–4 (approximately 12 days in culture), whereas late-passage pBUCs included passages 10, 15, and 18 (approximately 50 days in culture). Immortalized BUCs were analyzed at passage 7 (early; approximately 21 days in culture) and passage 55 (late; approximately 165 days in culture). The passage groups were derived from the same respective cell populations maintained in culture for different durations. Culture duration was calculated from the time of initial cell isolation for pBUCs and from the time of immortalization for iBUCs. The numerical labels used for the BUC experimental groups (12, 50, 21, and 165) refer to approximate days in culture rather than passage numbers.

A total of six bovine donors contributed to the RNA-seq datasets. Biological replicates generally represented three independent culture experiments performed at different times using cells from the same donor. The exception was the early-passage pBUC group, in which the three biological replicates originated from three different cows. Despite this donor variation, these samples showed high transcriptional similarity, comparable to the within-condition similarity observed among independently cultured replicates from the same donor, such as pBAT. Detailed sample information, including donor origin, culture conditions, and actual passage numbers, is provided in Supplementary Table S1.

### RNA extraction and quality assessment

Total RNA was extracted using the PureLink RNA Mini Kit (Invitrogen, catalog no. 12183018A) according to the manufacturer’s instructions. For scaffold-cultured cells, scaffolds were washed three times with phosphate-buffered saline for 5 min, transferred to lysis buffer supplemented with β-mercaptoethanol, and mechanically homogenized using a sterile pestle. Samples were centrifuged for 3 min at 10,000 × g, and the supernatant was mixed with an equal volume of 70% ethanol before RNA purification according to the manufacturer’s protocol.

RNA concentration was measured using a NanoDrop ND-1000 spectrophotometer, and RNA integrity was assessed using an Agilent 2100 Bioanalyzer. All samples had RNA integrity number (RIN) values ≥6.5, with the majority exceeding 9.

### Library preparation and sequencing

Bulk MARS-seq 3′ RNA-seq library preparation and sequencing were performed at the Nancy and Stephen Grand Israel National Center for Personalized Medicine. Single-end 100-bp reads were generated using an Illumina NovaSeq SP platform, with approximately 25 million reads obtained per sample.

### Bioinformatic processing

Initial bioinformatic processing was performed at the Mantoux Bioinformatics Institute, Weizmann Institute of Science. Poly-A/T stretches and Illumina adapter sequences were removed using cutadapt and reads shorter than 30 bp after trimming were discarded. Remaining reads were aligned using STAR in EndToEnd mode to 1,000-bp regions corresponding to annotated 3′ gene ends in the Bos taurus ARS-UCD1.2 reference genome based on RefSeq annotations, with outFilterMismatchNoverLmax = 0.05.

Reads mapping to the same gene and carrying identical unique molecular identifiers (UMIs) were collapsed to remove PCR duplicates. Gene-level quantification was performed using HTSeq-count, and UMI counts were corrected for saturation based on the expected number of unique molecules under sampling without replacement. The pipeline was implemented using Snakemake. The resulting gene-level UMI count matrix was used for downstream statistical analyses.

### Differential expression and downstream transcriptomic analyses

Genes with a total count ≤10 across the samples included in each analysis were excluded. Normalization and differential expression analysis were performed using DESeq2. Differentially expressed genes (DEGs) were defined as genes with a Benjamini–Hochberg adjusted p-value <0.05 and an absolute log2 fold change ≥1, unless specified otherwise.

Differential expression was assessed using both pairwise and one-versus-all comparisons. One-versus-all differential expression was assessed using DESeq2, comparing each condition with the equally weighted mean of all remaining conditions.

DESeq2-transformed expression values were used for PCA, PERMANOVA, correlation analyses, and expression-pattern clustering. For k-means clustering, expression values were additionally standardized by gene-wise z-score. The number of clusters was evaluated using the elbow method.

Functional enrichment analyses of DEG sets obtained from both pairwise and one-versus-all comparisons were performed using g:Profiler through the gprofiler2 R package. The statistical background was defined as the corresponding set of expressed genes that passed filtering and were tested in each analysis. Default significance criteria were applied, and only enriched terms containing fewer than 350 genes were retained unless specified otherwise. Gene Ontology Biological Process (GO:BP) and KEGG enrichment profiles obtained from one-versus-all analyses were additionally compared between groups using Jaccard similarity matrices based on overlap among significantly enriched terms.

Gene set enrichment analysis (GSEA v4.3.2) was performed using all tested genes ranked by the DESeq2 Wald statistic (stat). Normalized enrichment scores (NES) were used to compare pathway enrichment across predefined experimental contrasts.

KEGG pathway expression was visualized using the R package pathview (Luo and Brouwer, 2013). Genes with adjusted p < 0.05 were colored red (upregulated) or blue (downregulated). The same thresholds were used for the pathway enrichment shown in Figure 5.

Where indicated, enriched Gene Ontology terms were summarized using REVIGO (http://revigo.irb.hr/) to reduce semantic redundancy. Transcription factors identified through one-versus-all analyses were further examined using STRING v12.5 for network analysis.

PERMANOVA was used to assess associations between experimental factors and global transcriptomic variation using Euclidean distances calculated from DESeq2-transformed expression values. Tested factors included cell source, cell state, culture architecture, myogenic stage, passage group, and other comparison-specific variables, as appropriate. Gene-level R² values were calculated using linear models fitted independently for each gene to estimate the proportion of expression variance associated with the relevant experimental factor.

### Availability of data and materials

The RNA-seq datasets analyzed in this study are available in the Gene Expression Omnibus under accession numbers GSE276658, GSE327833, and GSE329893. GSE276658 and GSE327833 contain subsets previously deposited for related studies, whereas GSE329893 contains the additional samples required to complete the combined dataset analyzed here.

For reviewer access to GSE329893 prior to public release, use the token: **slaxwwkgpludfex**.

