## supplemental figures for "Comparative transcriptomics reveals shared and context-dependent adaptations of bovine cells across culture systems for cultivated meat"

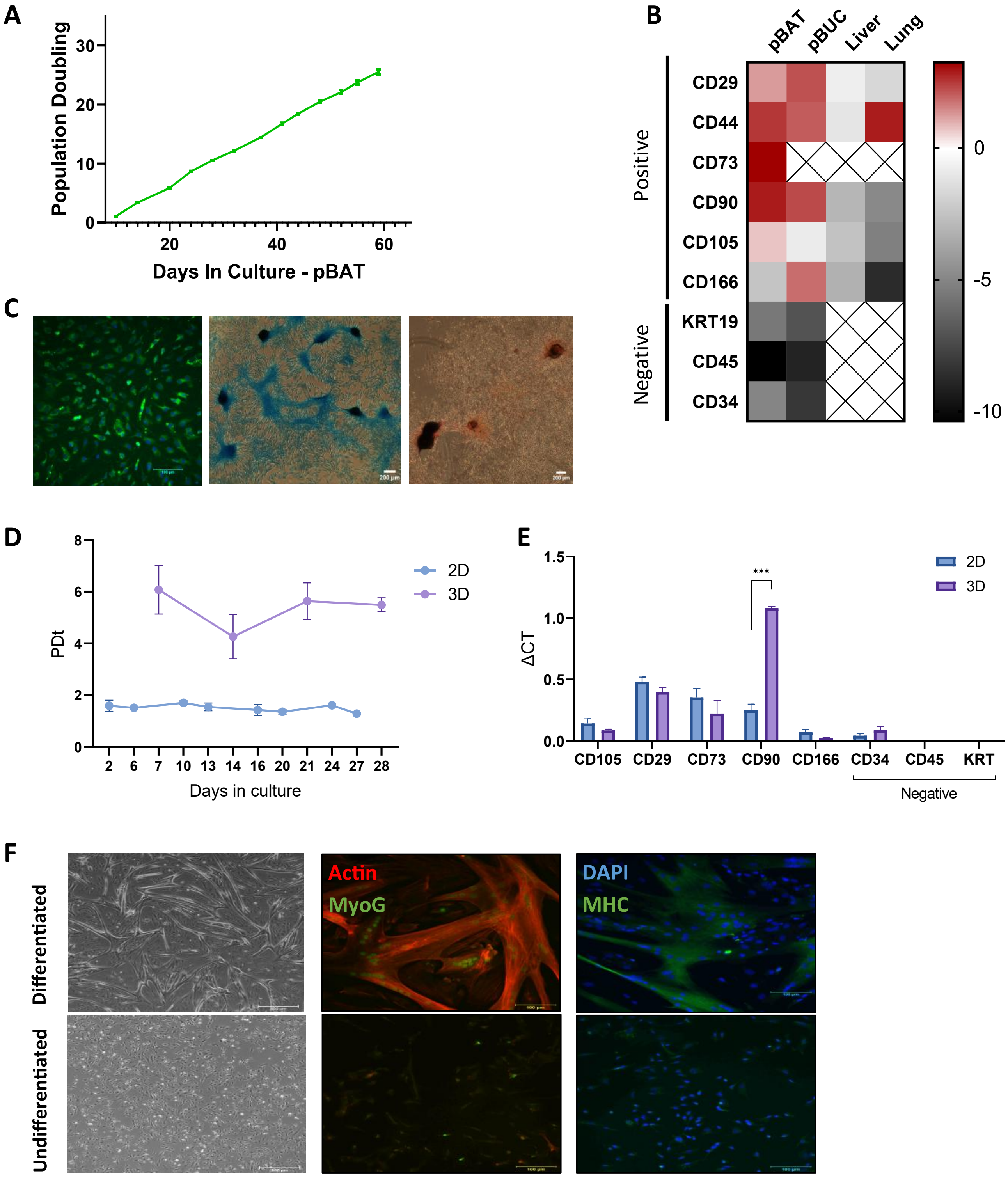
A
Population Doubling
Days In Culture - pBAT
B
pBAT
pBUC
Liver
Lung
Positive
Negative
CD29
CD44
CD73
CD90
CD105
CD166
KRT19
CD45
CD34
0
-5
-10
C
D
PDt
Days in culture
2D
3D
E
ΔCT
***
CD105
CD29
CD73
CD90
CD166
CD34
CD45
KRT
Negative
F
Differentiated
Undifferentiated
Actin
MyoG
DAPI
MHC

***Supplementary Figure S1. Characterization of the primary cell populations and spheroid cultures used in this study**. **A-C.** Characterization of primary bovine adipose tissue-derived cells (pBAT). **A.** Population doubling time, 3-4 biological replicates at each time point. PDt = SQRT(N/N0)/t (where SQRT, N, N0, and t are the square root, final cell numbers, the initial cell number, and the cell time in the culture, respectively). **B.** Expression of positive (CD29, CD44, CD73, CD90, CD105 and CD166) and negative (KRT19, CD34, CD45) MSC markers normalized to PSMB. **C.** Representative images showing adipogenic, chondrogenic, and osteogenic differentiation of pBAT, visualized by BODIPY 493/503 and DAPI staining, Alcian blue staining, and Alizarin red staining, respectively. Images were acquired using an EVOS FL Auto microscope. Scale bars, 100 µm. **D-E.** Characterization of pBAT spheroid cultures. **D.** Population doubling time of cells cultured in two-dimensional (2D) and three-dimensional (3D) conditions over time. Data are presented as mean ± SEM (n=3). **E.** Relative expression of positive and negative mesenchymal stromal cell marker genes in 2D and 3D cultures, measured by RT–qPCR and normalized to PSMB. Data are presented as mean ± SEM (n = 3 for 2D and n = 2 for 3D). *** p<0.001. **F.** Characterization of primary bovine satellite cells (pSAT). Representative bright-field and immunofluorescence images of undifferentiated and day-3 differentiated pSAT. Cells were stained for myogenin (MyoG; green) or myosin heavy chain (MHC; green), together with phalloidin-labelled actin (red) and DAPI (blue). Scale bars, 500 µm for bright-field images and 100 µm for immunofluorescence images.*

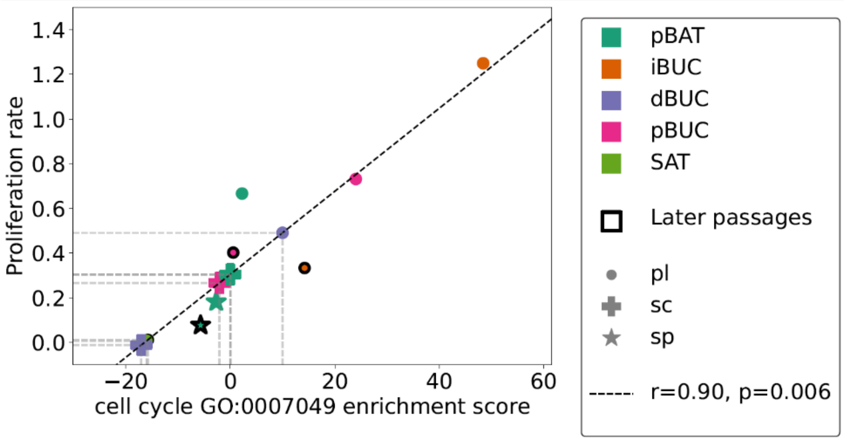


***Supplementary Figure S2.** Relationship between cell-cycle biological process enrichment and measured proliferation rates.*

A

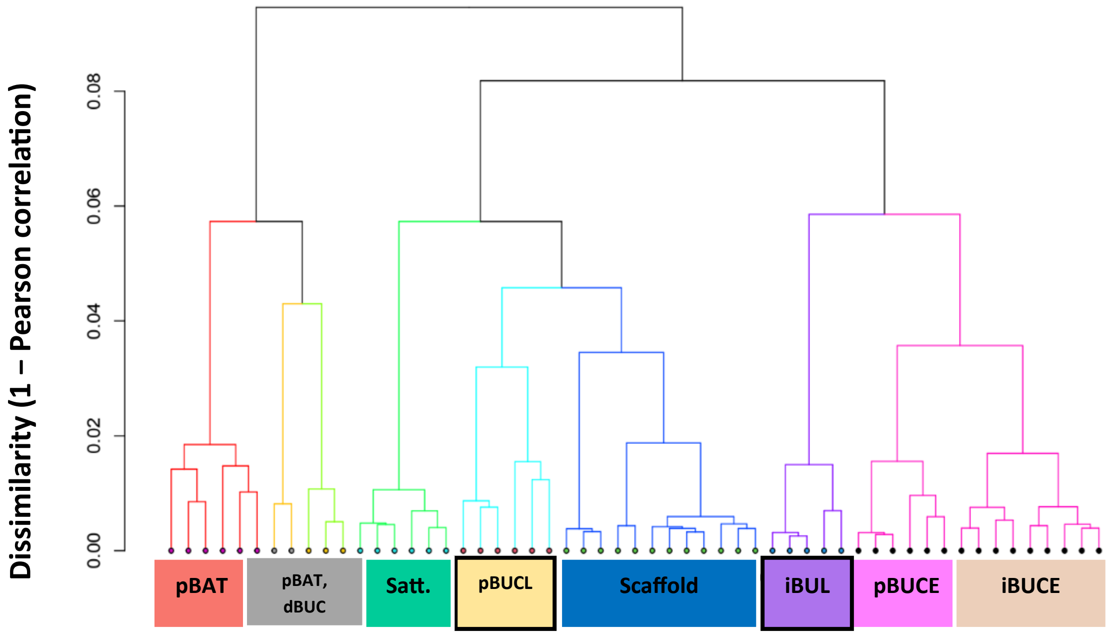
Dissimilarity (1 – Pearson correlation)
0.00
0.02
0.04
0.06
0.08
pBAT
pBAT, dBUC
Satt.
pBUCL
Scaffold
iBUL
pBUCE
iBUCE

B

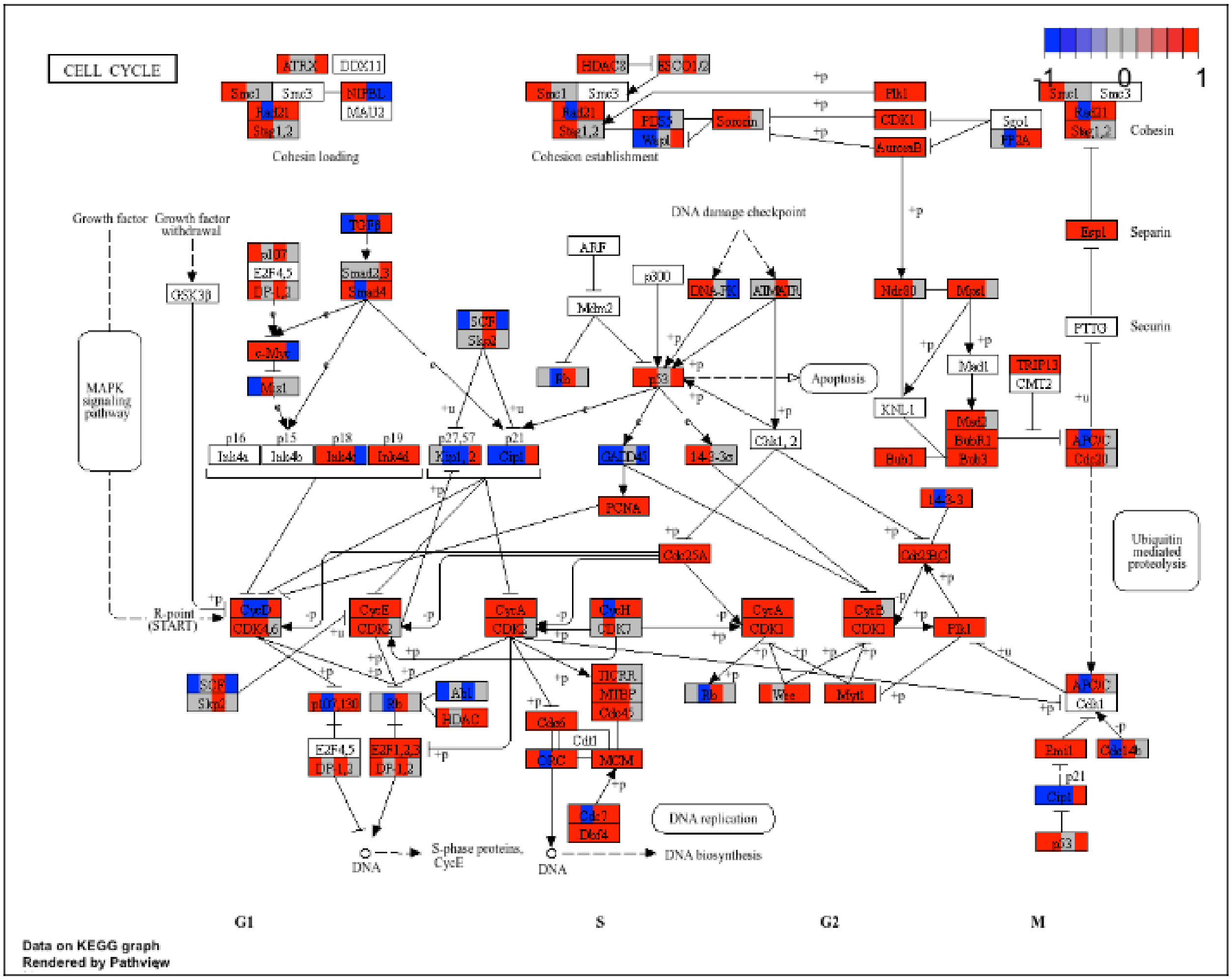
CELL CYCLE
-1
0
1
Cohesin loading
Cohesion establishment
Cohesin
Separin
Securin
Growth factor
Growth factor withdrawal
MAPK signaling pathway
DNA damage checkpoint
Apoptosis
Ubiquitin mediated proteolysis
R-point (START)
DNA replication
S-phase proteins, CycE
DNA
DNA biosynthesis
G1
S
G2
M
Data on KEGG graph
Rendered by Pathview

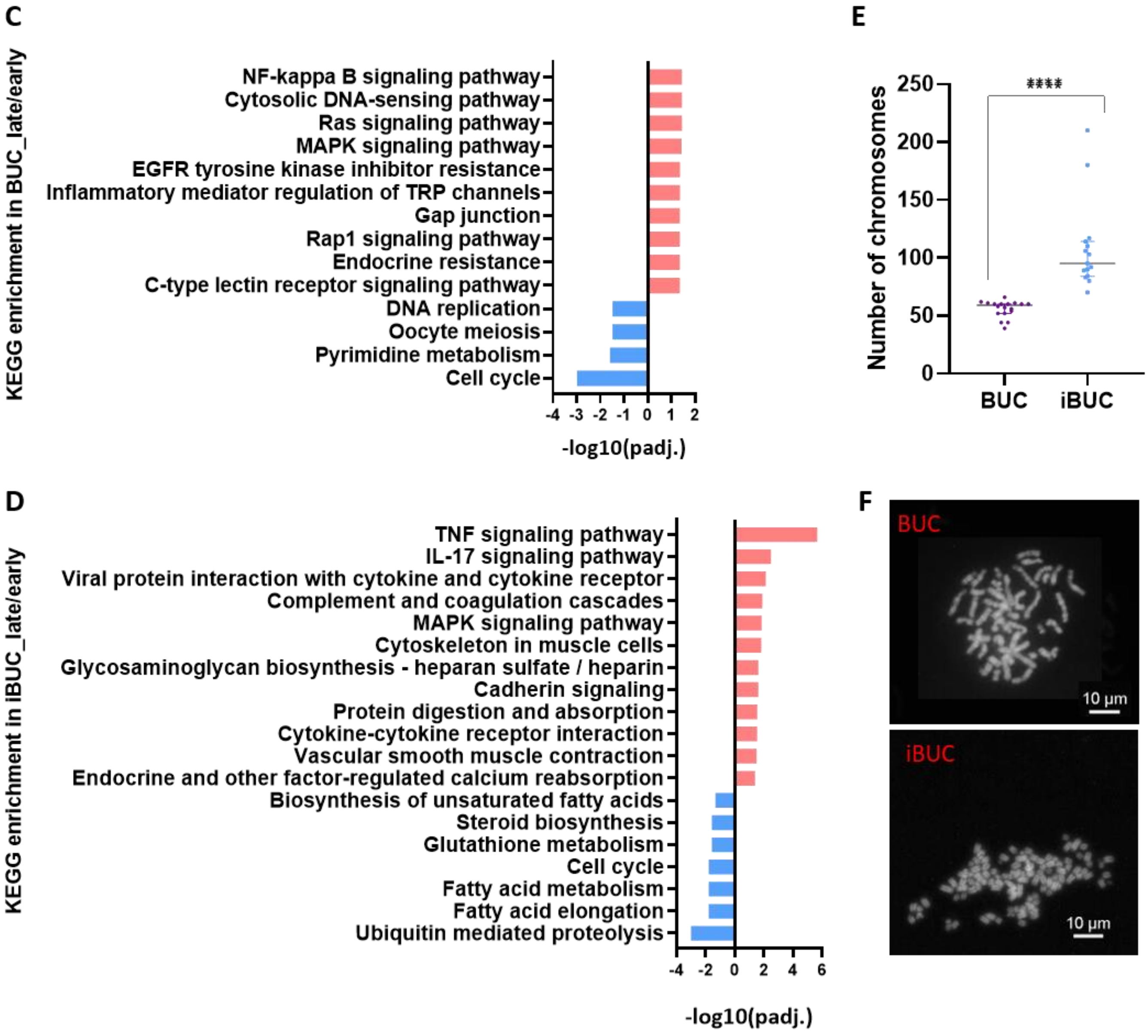


***Supplementary Figure S3. Transcriptomic relationships and pathway-level changes associated with BUC aging and immortalization.*** ***A.*** *Hierarchical clustering of samples by cell identity/state, culture architecture, and passage using complete linkage. The positions of early- and late-passage primary and immortalized BUC populations are indicated.* ***B.*** *Cell-cycle gene expression in early- and late-passage primary and immortalized BUC populations (adjusted p < 0.05).* ***C.*** *KEGG enrichment in BUC late/early.* ***D.*** *KEGG enrichment in iBUC late/early.* ***E.*** *Quantification of chromosome numbers per metaphase in BUC and iBUC cells, revealing aneuploidy in immortalized cells. Median ± 95% CI; BUC n = 18, iBUC n = 15 metaphases. Mann–Whitney test; significance is marked by ****p < 0.0001.* ***F.*** *Representative metaphase chromosome spreads of primary BUC and iBUC cells. Scale bars, 10 µm.*

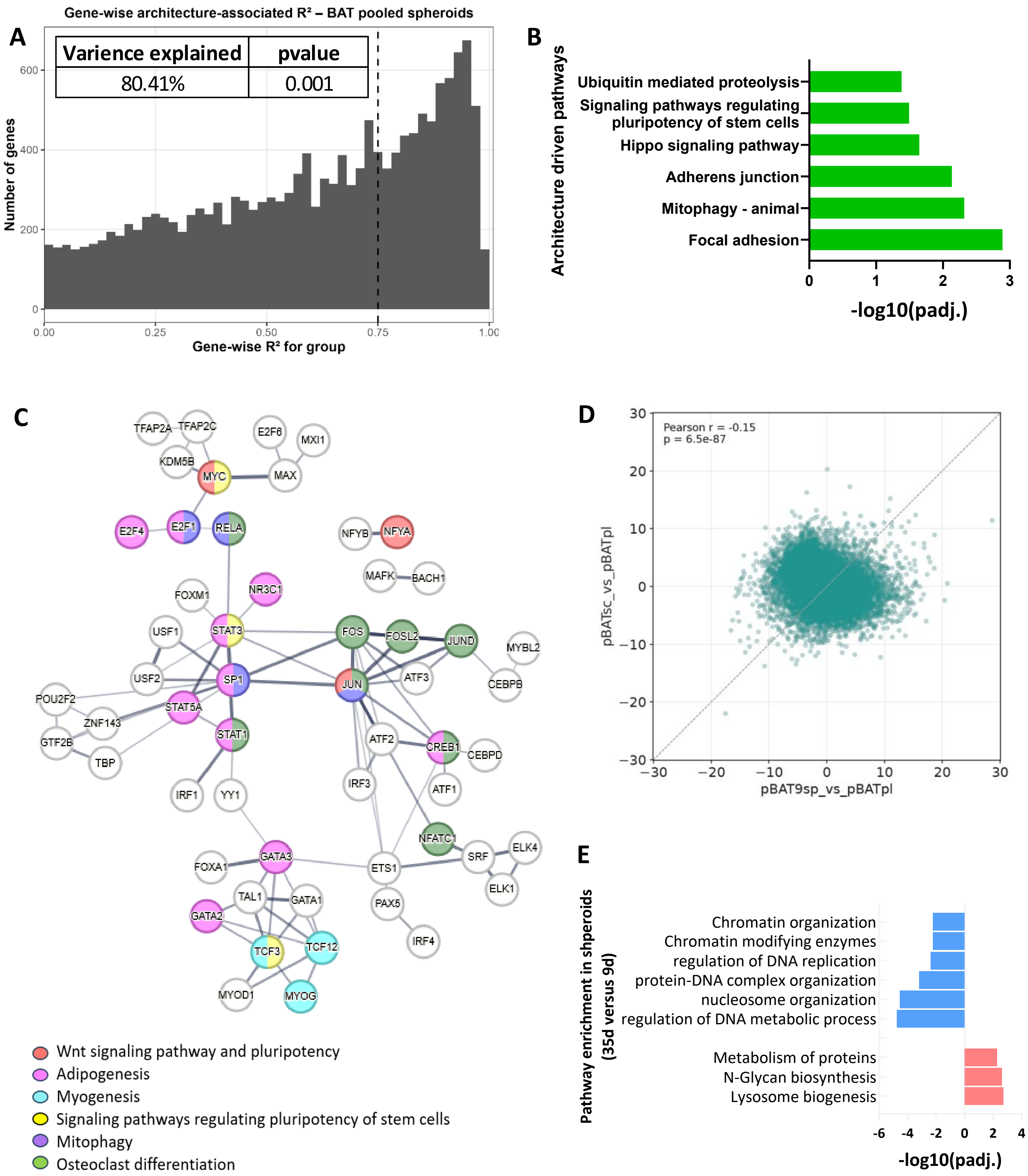
A
Gene-wise architecture-associated R² – BAT pooled spheroids
Varience explained
pvalue
80.41%
0.001
Number of genes
Gene-wise R² for group
0
200
400
600
0.00
0.25
0.50
0.75
1.00
B
Architecture driven pathways
Ubiquitin mediated proteolysis
Signaling pathways regulating pluripotency of stem cells
Hippo signaling pathway
Adherens junction
Mitophagy - animal
Focal adhesion
0
1
2
3
-log10(padj.)
C
TFAP2A
TFAP2C
E2F6
MXI1
KDM5B
MYC
MAX
E2F4
E2F1
RELA
NFYB
NFYA
MAFK
BACH1
FOXM1
NR3C1
USF1
STAT3
FOS
FOSL2
JUND
MYBL2
USF2
SP1
JUN
ATF3
CEBPB
POU2F2
STAT5A
ZNF143
GTF2B
STAT1
ATF2
CREB1
CEBPD
TBP
IRF1
YY1
IRF3
ATF1
NFATC1
SRF
ELK4
FOXA1
GATA3
ETS1
ELK1
TAL1
GATA1
PAX5
GATA2
TCF3
TCF12
IRF4
MYOD1
MYOG
Wnt signaling pathway and pluripotency
Adipogenesis
Myogenesis
Signaling pathways regulating pluripotency of stem cells
Mitophagy
Osteoclast differentiation
D
Pearson r = -0.15
p = 6.5e-87
pBATsc_vs_pBATpl
pBAT9sp_vs_pBATpl
−30
−20
−10
0
10
20
30
E
Pathway enrichment in shperoids (35d versus 9d)
Chromatin organization
Chromatin modifying enzymes
regulation of DNA replication
protein-DNA complex organization
nucleosome organization
regulation of DNA metabolic process
Metabolism of proteins
N-Glycan biosynthesis
Lysosome biogenesis
-6
-4
-2
0
2
4
-log10(padj.)

***Supplementary Figure S4. Architecture-associated transcriptional and pathway changes in pBAT.*** ***A.*** *Distribution of gene-wise architecture-associated $R^2$ values in pBAT, with the overall architecture-associated variance and significance indicated.* ***B.*** *Selected KEGG pathways associated with culture architecture. Bars represent −log10(adjusted p- value).* ***C.*** *Transcription-factor interaction network associated with spheroid culture, highlighting downregulated transcription factors linked to developmental and differentiation-related programs.* ***D****. Gene-wise DESeq2 Wald statistics for scaffold-versus-plate vs spheroid-versus-plate contrasts, with Pearson correlation coefficient.* ***E****. Biological Process enrichment associated with prolonged spheroid culture (35-day versus 9-day spheroids), showing processes enriched among up- and downregulated genes.*

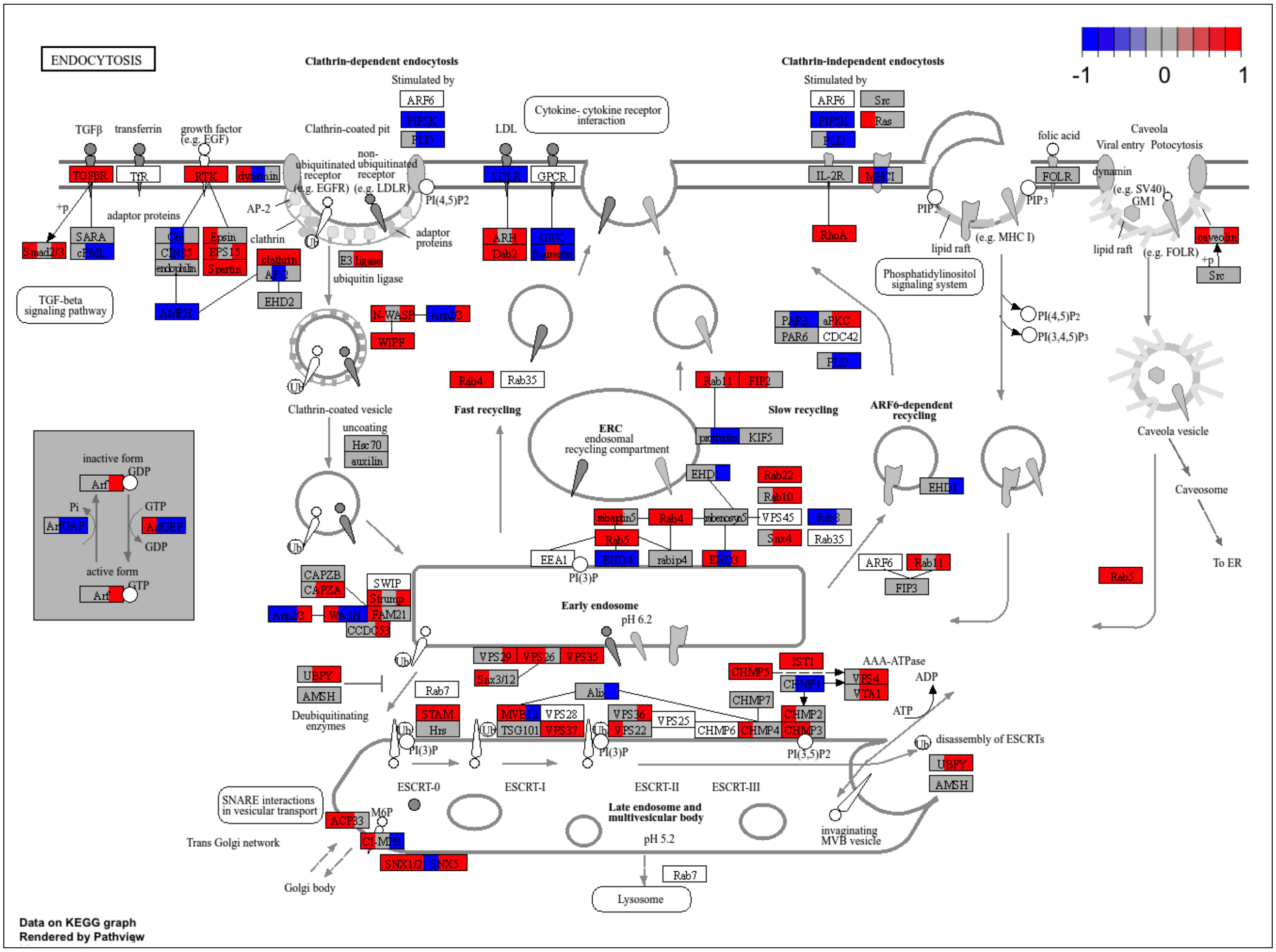


***Supplementary Figure S5. Differential regulation of the endocytosis pathway in scaffold versus plate culture.*** *KEGG endocytosis pathway genes are colored according to differential expression in three scaffold-versus-plate comparisons: pBUC (left), pBAT (center), and dBUC (right). Red indicates significantly higher expression on scaffold and blue indicates significantly lower expression on scaffold (adjusted $p < 0.05$).*

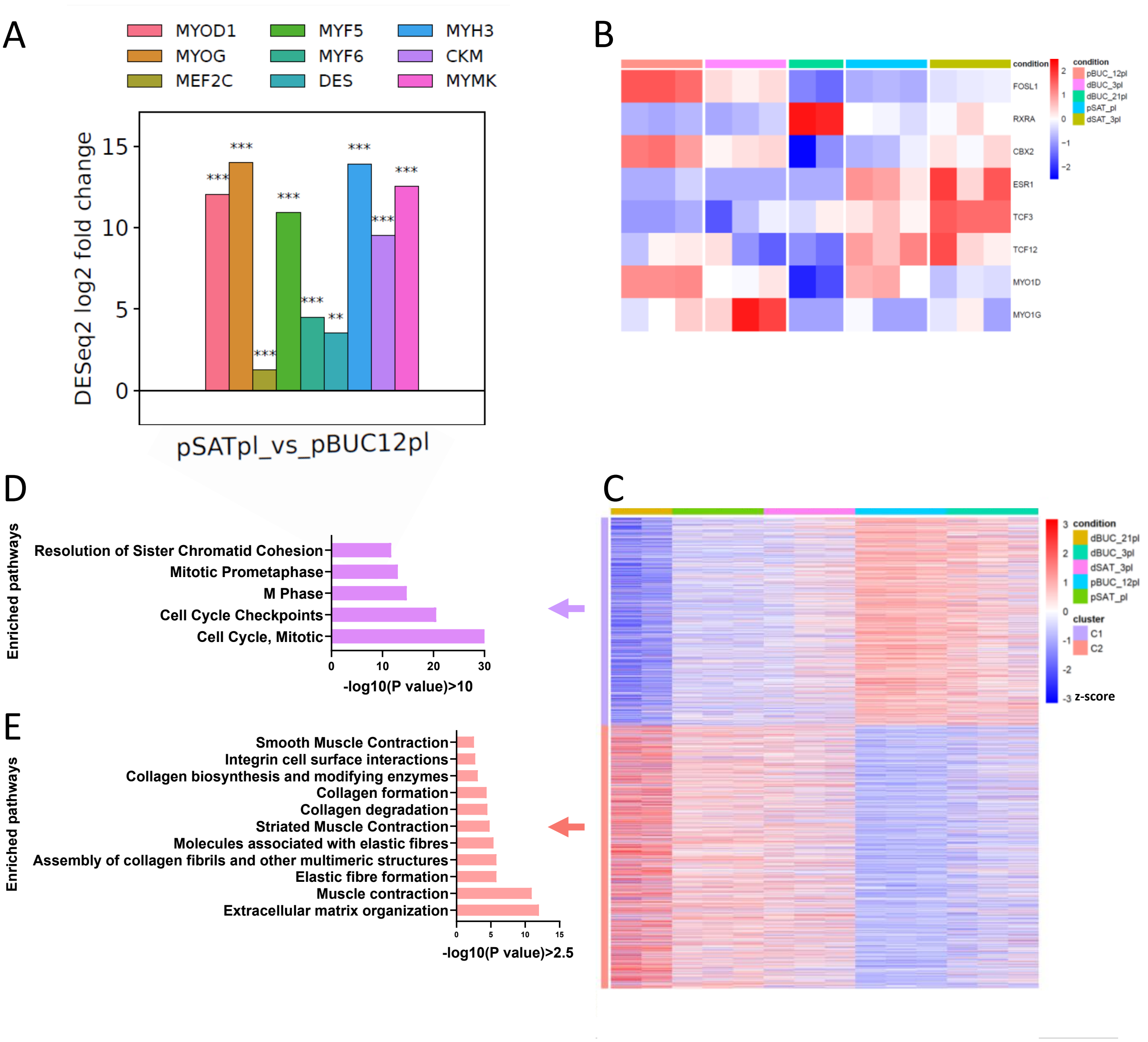


***Supplementary Figure S6. Detailed transcriptional programs associated with BUC and satellite-cell myogenesis.***

***A.*** *Expression of selected myogenic genes in undifferentiated satellite cells and BUCs.* ***B.*** *Transcription-factor profiles of myogenic cell populations identified by one-versus-all comparison.* ***C.*** *Clustering of differentiation-associated genes across plate-cultured BUC and satellite-cell populations.* ***D-E****. Pathway enrichment of clusters in C.*
